# EspH utilizes phosphoinositide and Rab binding domains to interact with plasma membrane infection sites and Rab GTPases

**DOI:** 10.1101/2024.06.28.601186

**Authors:** Ipsita Nandi, Rachana Pattani Ramachandran, Deborah E. Shalev, Dina Schneidman-Duhovny, Raisa Shtuhin-Rahav, Naomi Melamed-Book, Ilan Rosenshine, Benjamin Aroeti

## Abstract

Enteropathogenic *E. coli* (EPEC) is a Gram-negative bacterial pathogen that causes persistent diarrhea. Upon attachment to the apical plasma membrane of the intestinal epithelium, the pathogen translocates virulent proteins called effectors into the infected cells. These effectors hijack numerous host processes for the pathogen’s benefit. Therefore, studying the mechanisms underlying their action is crucial for a better understanding of the disease. We show that translocated EspH interacts with multiple host Rab GTPases. AlphaFold predictions and site-directed mutagenesis identified glutamic acid and lysine at positions 37 and 41 as Rab interacting residues. Mutating these sites abolished the EspH ability to inhibit the Akt and mTORC1 signaling, lysosomal exocytosis, and bacterial invasion. Knocking out the endogenous Rab8a gene expression highlighted the involvement of Rab8a in Akt/mTORC1 signaling and lysosomal exocytosis. A phosphoinositide binding domain with a critical tyrosine was identified in EspH. Mutating the tyrosine abolished the localization of EspH at infection sites and its capacity to interact with Rabs. Our data suggest novel EspH-dependent mechanisms that elicit immune signaling and membrane trafficking during EPEC infection.

## Introduction

The human gastrointestinal pathogens, enteropathogenic and enterohemorrhagic *Escherichia coli* (EPEC and EHEC), are food or water-borne bacterial pathogens that continue to pose a significant threat to human health worldwide (Kaur & Dudeja, 2023; Nataro & Kaper, 1998; Ochoa & Contreras, 2011). *Citrobacter rodentium* is a natural murine intestinal pathogen that shares a set of virulence factors with EPEC and EHEC (Silberger et al, 2017). The type III secretion system (T3SS), which has a syringe-like molecular structure, is a major virulence factor of these pathogens (Slater et al, 2018). They utilize it to inject dozens of proteins, termed effectors, from the bacterial cytoplasm into the host enterocytes. The coordinated activity of these effectors in space and time subverts numerous host cell processes and organelles to support successful bacterial colonization of the intestinal mucosa (Sanchez-Garrido et al, 2021). A prominent hallmark of the infection is the appearance of the so-called “attaching and effacing” (A/E) lesions in the mucosal tissue. A/E lesions are characterized by intimate microbial attachment to the apical cell plasma membrane of the epithelial cells, local elimination of brush-border microvilli, and the formation of a filamentous (F)-actin-rich pedestal-like structure on top of which the bacterium resides. Studies suggest that the A/E pedestal formation contributes to bacterial pathogenesis (Miner & Rauch, 2024). Injected effectors also contribute to bacterial colonization and pathogenesis by hijacking and subverting host cell signaling and membrane trafficking pathways, targeting various organelles, and modulating cell death and innate immunity (El Qaidi et al, 2019; Lee et al, 2024; Shenoy et al, 2018). One such effector is EspH.

EspH is essential for virulence and is a multifunctional effector (Ritchie & Waldor, 2005; Wong et al, 2011). It regulates the actin cytoskeleton and pedestal formation (Wong et al, 2012b), conferring Rho GTPase (Dong et al, 2010a; Dong et al, 2010b; Ramachandran et al, 2022; Roxas et al, 2018; Tu et al, 2003; Wong et al, 2012a) and MAP kinase (Ramachandran et al, 2018) inhibition, inducing host cell cytotoxicity (Ramachandran et al, 2022; Tu et al, 2003), inhibiting bacterial invasion (phagocytosis) (Dong et al, 2010b; Ramachandran et al, 2022; Wong et al, 2011), and increasing cell death due to mitochondrial fragmentation (Roxas et al, 2022). Following translocation into the host cell, EspH localizes at plasma membrane infection sites (Ramachandran et al, 2022; Ramachandran et al, 2018; Tu et al, 2003). Despite this knowledge, the molecular mechanisms underlying EspH’s activities are not fully understood.

Rab GTPases constitute the largest subfamily of the Ras-related GTPase superfamily, regulating multiple steps in vesicular trafficking in eukaryotic cells (Herve & Bourmeyster, 2018; Hutagalung & Novick, 2011; Zerial & McBride, 2001). Like all members of the GTPase superfamily, Rabs can bind GDP (guanosine-5’-diphosphate) in the “off-state” or GTP (guanosine-5’-triphosphate) in the “on-state.” Host proteins that regulate cycling between the two states are guanine nucleotide exchange factors (GEFs) that catalyze the GDP-GTP exchange reaction and GTPase activating proteins (GAPs) that facilitate the hydrolysis of bound GTP, thereby switching the Rabs on and off, respectively (Novick, 2016), and targeting them to distinct cellular compartments (Pfeffer & Aivazian, 2004). Switching between the active and inactive forms is characterized by conformational changes of two critical regions within the GTPases, called switch I and II (Vetter & Wittinghofer, 2001). The Rab proteins also interact with an additional group of proteins called “effector” proteins. These proteins bind with high affinity to the switch regions of Rab’s active form (Gillingham et al, 2014; Khan, 2013), exerting specific cellular functions, some of which are overridden by intracellular bacterial pathogens (Spano & Galan, 2018; Stein et al, 2012).

Studies have shown that human bacterial pathogens (e.g., *Salmonella enterica*, *Legionella pneumophila*, *Shigella flexneri*, *Listeria monocytogenes*) target Rab GTPases to establish a replication niche that enables their survival within the host cells (Dubreuil & Segev, 2011; Sherwood & Roy, 2013; Stein et al, 2012). Moreover, studies have indicated that type III secreted effectors can target host Rab GTPases directly (Boddy et al, 2021; Gan et al, 2020; Goody et al, 2012; Mousnier et al, 2014; Spano et al, 2016). *Salmonella’s* SopD and *Legionella’s* LidA are, perhaps, the best-studied examples, as crystal structures with their Rab cognates have been resolved (Cheng et al, 2012; Lian et al, 2021; Schoebel et al, 2011). SopD binds and displays a GAP activity that inhibits Rab8a (Lian et al, 2021; Savitskiy & Itzen, 2021) and Rab10 (Boddy et al, 2021). Effector proteins can also act as GEFs. For example, the *Legionella’s* SidM (DrrA) effector recruits and activates Rab1 via its GEF domain. The same effector also acts as a GDP dissociation inhibitor displacement factor of Rab1 (Ingmundson et al, 2007; Machner & Isberg, 2007; Murata et al, 2006). Effector proteins can modulate the activity of Rabs by covalently modifying them. For example, the AMPylation activity of the *Legionella* SidM/DrrA modifies Rab1 by covalently adding adenosine monophosphate (AMP) to avoid its recognition by GAPs (Muller et al, 2010). Another effector protein, SidD, acts as deAMPylase (Neunuebel et al, 2011; Tan & Luo, 2011)]. Type III secreted effectors can also display protease activities on Rabs, such as *Salmonella*’s GtgE effector, which acts as a Rab32 protease (Spano et al, 2016).

The *salmonella* effector protein, SopD, has been shown to enhance or inhibit inflammatory responses by targeting Rab8a signaling (Lian et al, 2021). The *Legionella* LidA-Rab6a interactions were shown to be required for bacterial intracellular replication and growth (Chen & Machner, 2013). Notably, these effector-rab structure-function relationships have been characterized for invading bacterial pathogens. As far as we know, no knowledge exists about such relations for extracellular bacterial pathogens, including the A/E pathogens. Here, we show for the first time that translocated EspH functionally interacts with multiple active Rabs, including Rab8a, Rab10, Rab3a, and Rab12. We also show that EspH possesses a putative phosphoinositide binding domain (PBD), which plays a pivotal role in EspH localization at plasma membrane infection sites, likely by mediating effector binding to phosphoinositides (PIs). The PBD is also vital for maintaining the EspH-Rab interactions.

## Results

### Translocated EspH interacts with the human Rab GTPases Rab8a, 10, 3a, and 12

Using co-precipitation followed by mass spectrometry and proteomics analysis, we have previously shown that EspH co-precipitated with multiple Rab GTPases, including Rab8a, Rab10, Rab3(a and d), and Rab12, Rab1(a and b) and Rab39a. It has been suggested that these Rabs interact with a region in EspH located upstream of the C-terminal 38-amino acid (aa) segment (Ramachandran et al, 2022). The current study focused mainly on Rab8a, Rab10, Rab3a, and, to some extent, Rab12. To confirm these findings by immunoblotting (IB), HeLa or Caco-2_BBe_ cells were infected with an *espH* deleted EPEC strain (EPEC-Δ*espH*), or with an EPEC-Δ*espH* mutant complemented with a plasmid expressing wild-type (*wt*) EspH with six histidines (6xHis) and a streptavidin binding peptide (SBP) tags located at the C-terminus of the effector protein (EPEC-Δ*espH*/pEspH*_wt_*). Notably, these experiments used cell infection with EPEC-Δ*espH* as negative controls. Cells were lysed, and EspH was precipitated using Streptavidin (StAv) agarose beads. SDS-PAGE, followed by IB, was used to analyze precipitated EspH and co-precipitated Rabs. The results showed that the Rabs co-precipitated with EspH from the EPEC-Δ*espH*/pEspH*_wt_* infected cells (**Fig. 1**). Additionally, the endogenous Rab8a co-precipitated with EspH from EPEC-Δ*espH*/pEspH*_wt_* and from cells infected with EPEC-Δ*espH* complemented with a plasmid expressing EspH whose C-terminal 38 aa ABR binding domain (**Fig. S1)** was deleted (EPEC-Δ*espH*/pEspH*_Δ130-168_*; **Fig. S2**). These results are consistent with our proteomics analyses (Ramachandran et al, 2022), suggesting that the Rabs interact with the translocated EspH via a region located upstream to the C-terminal 38 aa segment.

**Fig. 1:**
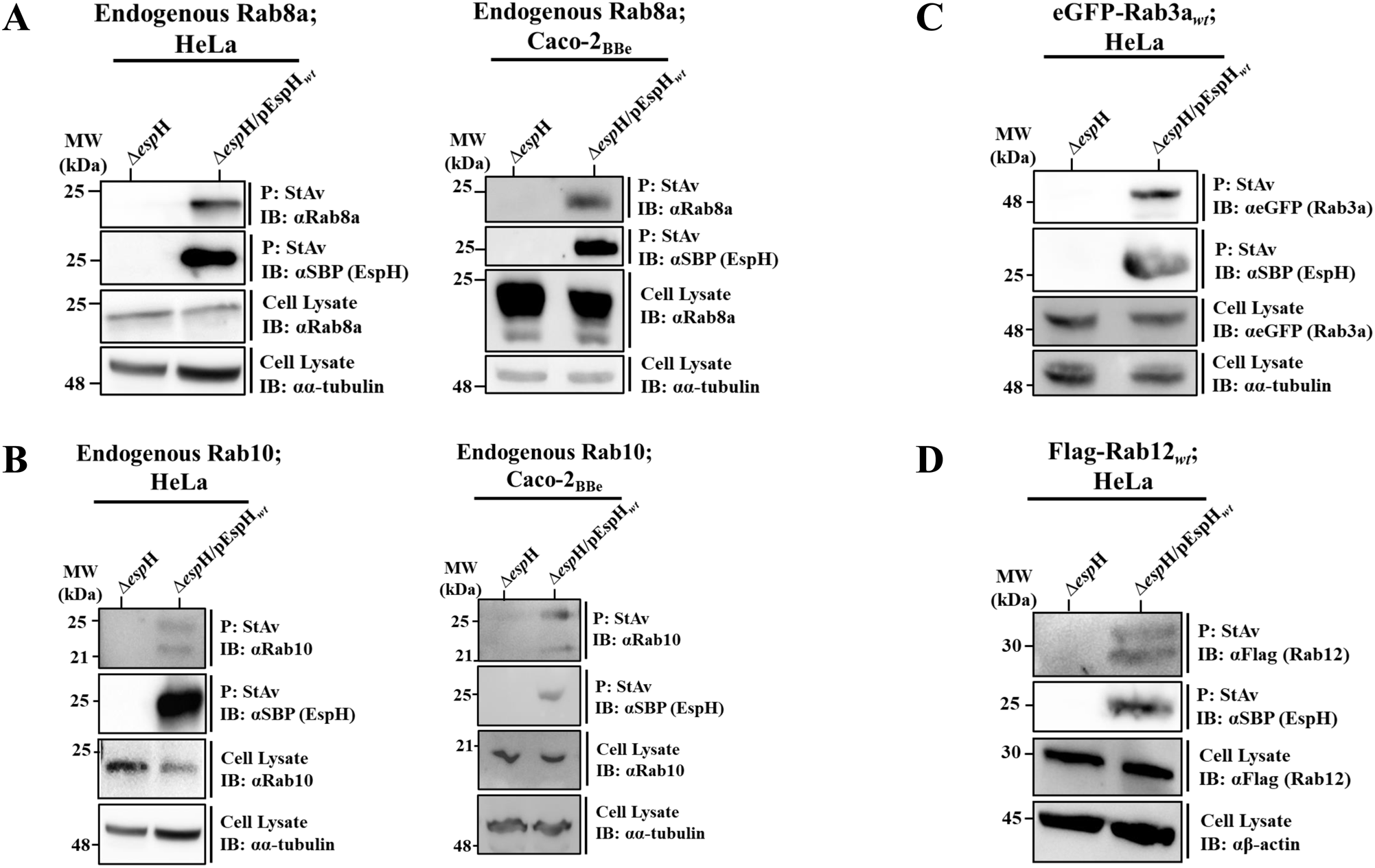
Rab8a, Rab10, Rab3a and Rab12 co-precipitate with translocated EspH*_wt_*. HeLa cells were infected with the indicated EPEC strains, lysed, and subjected to co-precipitation analyses, as described in Materials and Methods. SBP-tagged EspH*_wt_* was precipitated (P) by streptavidin (StAv) beads, and anti-SBP antibodies were used to identify the precipitated EspH by immunoblotting (IB). The co-precipitated endogenous Rab proteins were detected by IB, using antibodies directed against them (**panels A and B**) or anti-eGFP or anti-Flag tag antibodies in cases where epitope-tagged Rabs were ectopically expressed **(C and D**). The same antibodies were used to identify the Rabs in cell lysates. Anti-β-actin (αβ-actin) or anti-α-tubulin (αα-tubulin) antibodies were used for detecting protein loading. The endogenous Rab8a and Rab10 co-precipitation was examined in HeLa (**A**) and Caco-2_BBe_ (**B**) cells. Representative gels out of at least three independent experiments are shown.

### EspH interacts with the active Rab forms

Using the same co-precipitation approach, we next examined the capacity of translocated EspH*_wt_* to co-precipitate the ectopically expressed GFP-Rab8a*_wt_*, eGFP-Rab3a*_wt_* and eGFP-Rab10*_wt_*, the dominant negative (GDP-locked) GFP-Rab8a_T22N_; eGFP-Rab3a_T36N_; eGFP-Rab10_T23N_ and the constitutively active (GTP-locked) GFP-tagged Rab8a_Q67L_; eGFP-Rab3a_Q81L_; eGFP-Rab10_Q58L_. The *wt*, the constitutively active but not the dominant negative Rabs, co-precipitated with EspH*_wt_* from EPEC-Δ*espH*/pEspH*_wt_* infected cells (**Fig. 2 and Figs. S3A&S3B, left panels**). Confocal imaging confirmed these results, showing that translocated EspH*_wt_* colocalized with the *wt* and the constitutively active but not with the dominant negative Rabs (**Fig. 2 and Fig. S3A and S3B, middle and right panels**). These results suggest that EspH selectively binds active Rabs.

**Fig. 2:**
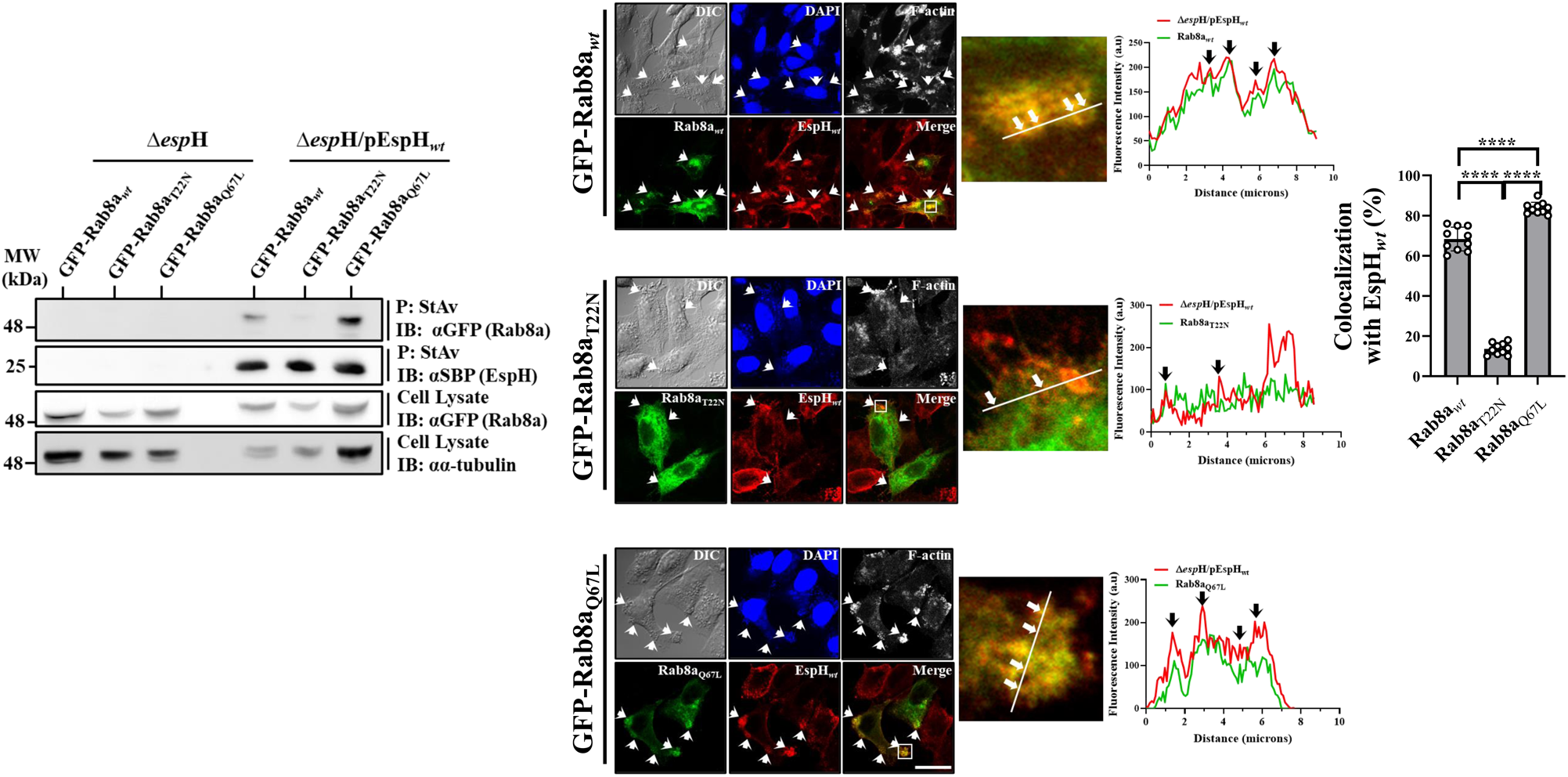
Translocated EspH*_wt_* interacts with active Rab8a. **Analysis by co-precipitation (left).** HeLa cells ectopically expressing the wild-type (*wt*), T22N (GDP-locked), and Q67L (GTP-locked) GFP-tagged Rab8a were infected with EPEC-Δ*esp*H or EPEC-Δ*esp*H/pEspH*_wt_* for 90 min at 37 °C and subjected to co-precipitation experiments, as in Fig. 1. A representative gel out of three independent experiments is shown. **Colocalization analysis (right).** HeLa cells expressing the eGFP-tagged Rabs were infected with EPEC-Δ*esp*H/pEspH*_wt_*, fixed, permeabilized, and immunostained with anti-SBP antibodies to visualize EspH. Cells were also stained with DAPI (to visualize cell nuclei and bacterial microcolonies) and Phalloidin CF-647 (F-actin). Cells were imaged by confocal microscopy, and representative images (out of 3 independent experiments) are shown. The green (Rab) and the red (EspH) channels were merged. Arrows point towards infecting EPEC microcolonies. An enlargement of the boxed region in the merged image is shown, along with a line used to generate fluorescence intensity profiles of EspH and Rab. Arrows point towards co-peaking fluorescence intensity signals. The bar graph depicts a quantitative colocalization analysis between the Rabs and EspH. Results are mean ± SE. The results of parallel experiments involving eGFP-Rab3a and eGFP-Rab10 are shown in **Fig. S3A** and **Fig. S3B**, respectively.

### AlphaFold calculated structures show a shared binding interface between EspH and the Rabs, where EspH residues E37 and K41 were implicated in Rab-binding

AlphaFold-Multimer-v2.0 was used to calculate complexes of EspH (pink) and the different Rabs (gray) (**Fig. 3A and 3B**). The <u>p</u>redicted <u>a</u>lignment <u>e</u>rror (PAE) showed high confidence in the binding regions of Rab3a and Rab10, with a somewhat reduced confidence in binding Rab8a and a significantly reduced confidence in binding Rab12 (**Fig. 3C**), suggesting that the interactions of EspH with these Rabs are weak. In all calculated structures, residues E37 and K41 located in an α-helix of EspH interacted with corresponding Lys (K) and Asp (D) residues located in a β-strand of the interswitch region of the Rabs. Specifically, E37 and K41 of EspH with D45 and K47 residues of Rab10 (**Fig. 3A**), D58 and K60 in Rab3a, D44 and K46 in Rab8a, and D78 and K80 in Rab12 (**Fig. 3B**). F10 in EspH seems to interact with Rab3a or Rab10 (**Fig. 3A and 3B**) via F59 and W76 in Rab3a and F46 and W63 in Rab10. Interestingly, the aromatic F and W residues in the Rabs are among the three amino acids that make up the hydrophobic triad defined as F45, W62, and Y77 of human Rab8a, which interacts with the Legionella LidA effector (Schoebel et al, 2011). These additional interactions correlate with the increased confidence of binding of Rab10 and Rab3a with EspH, shown in the PAE plots (**Fig. 3C**). Notably, the predicted Rab binding residues of EspH from different *E. coli* species are highly conserved, and as expected, are located upstream to the ABR binding C-terminal 38 aa domain (**Fig S1**).

**Fig. 3:**
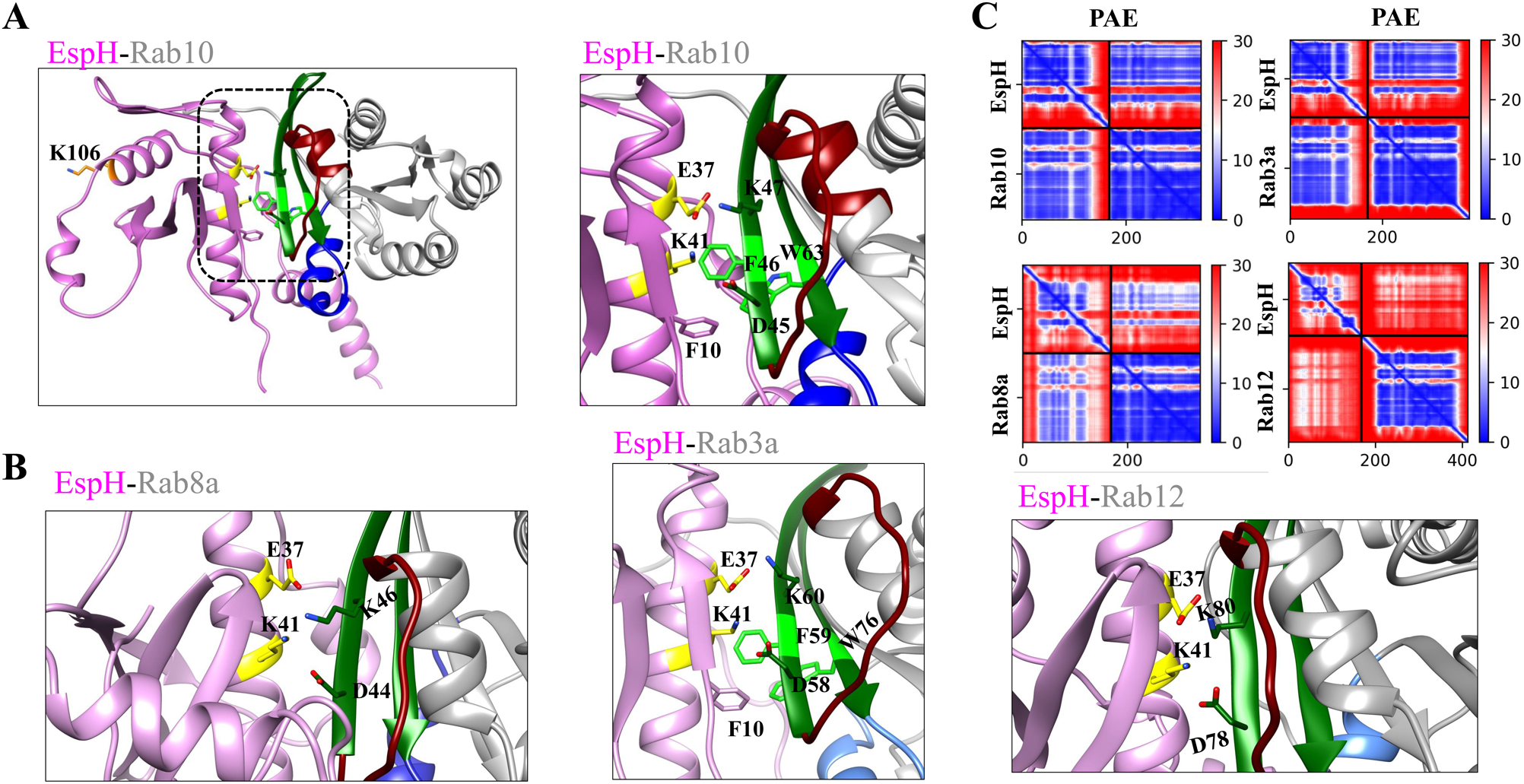
AlphaFold predicted structures for binding interfaces of EspH-Rabs. The complexes of Rab10-EspH, Rab8a-EspH, Rab3a-EspH, and Rab12-EspH were modeled using AlphaFold-Multimer-v2.0. The EspH (pink) and the Rab GTPase (gray) structures are depicted. The switch I, interswitch, and switch II Rab domains are shown in maroon, dark green, and navy blue, respectively. **(A). The EspH-Rab10 binding interface.** The Rab10-EspH complex is shown in the left panel, and the interface area (encircled) is enlarged in the right panel, where the predicted interacting E37 and K41 of EspH (yellow) and K47 and D45 of Rab10 interswitch region (green) are shown. Also, F10 (pink) of EspH interacts with F46 (bright green) and W63 (bright green) of Rab10. The non-interacting K106 of EspH (orange) is also indicated (left). **(B) The EspH and Rab8a (left), Rab3a (middle), and Rab12 (right) binding interfaces. (C) <u>P</u>redicted <u>A</u>ligned <u>E</u>rror (PAE) plots for EspH and the denoted Rab proteins.** The interacting E37 and K41 residues of EspH (yellow) respectively interact with K46 and D44 in Rab8A, with K60 and D58 in Rab3a, and with K80 and D78 in Rab12, all located in the interswitch region (green). F10 of EspH (pink) interacts with F59 and W76 in Rab3a.

### Residues E37 and K41 of EspH are crucial for Rab binding

The significance of E37 and K41 of EspH in Rab binding was investigated by mutating these residues individually to alanine (A) (**Fig. S1**). EPEC-Δ*espH* mutant strains complemented with plasmids encoding the EspH mutant EspH*_E37A_* (EPEC-Δ*espH*/pEspH*_E37A_*) or EspH*_K41A_*(EPEC-Δ*espH*/pEspH*_K41A_*) were generated, and their ability to translocate the corresponding EspH mutants was confirmed (**Fig. S4**). HeLa cells were infected with these bacterial strains and subjected to the Rab co-precipitation approach. The results show that EspH*_E37A_* and EspH*_K41A_* failed to co-precipitate endogenous Rab8a (**Fig. 4A**), eGFP-Rab3a*_wt_* (**Fig. 4B**), and the endogenous Rab10 (**Fig. 4C**). Similar results were obtained with the endogenous Rab8a in Caco-2_BBe_ infected cells (**Fig. S5**).

**Fig. 4:**
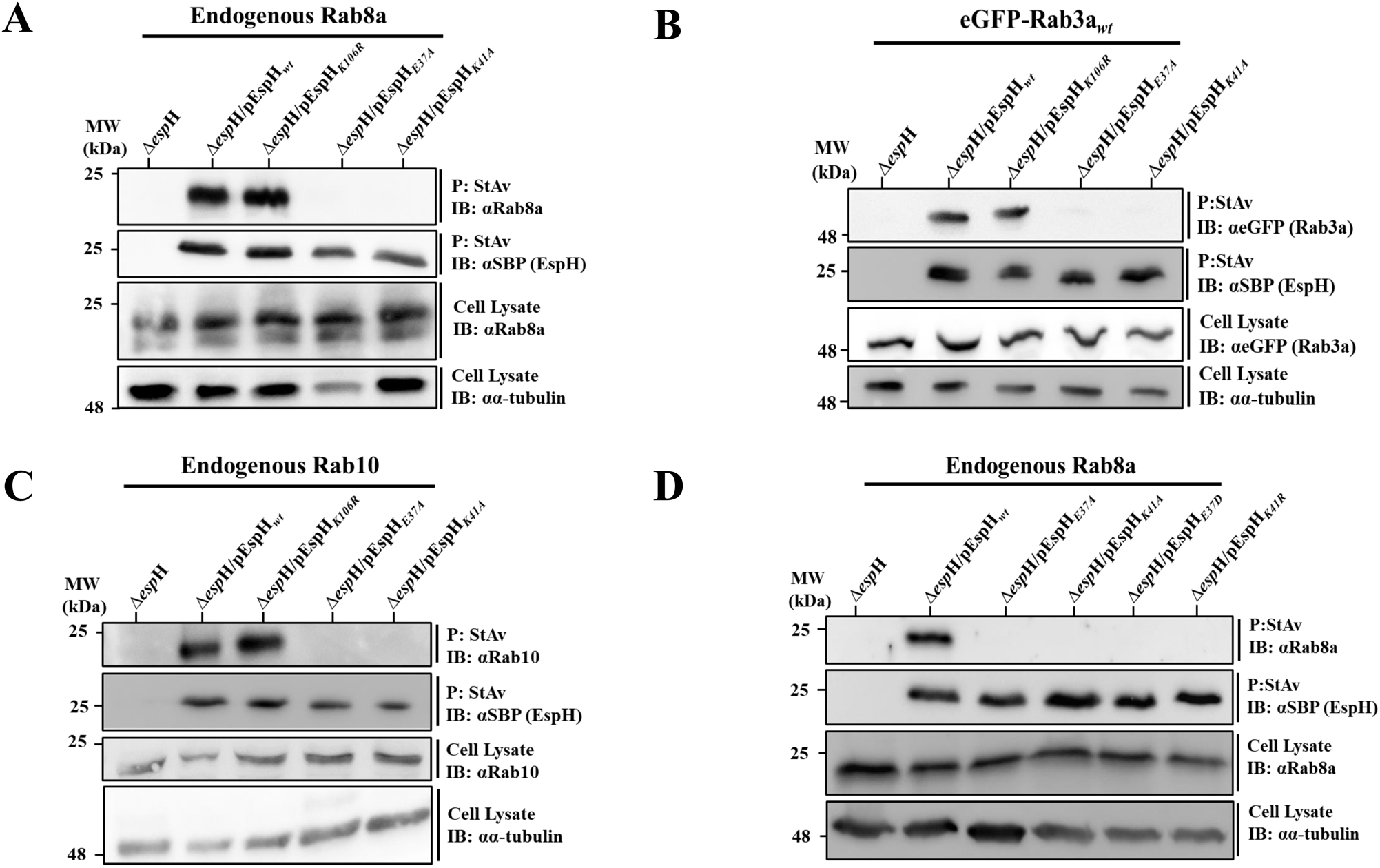
The predicted Rab binding residues in EspH are critical for EspH-Rab interactions and are highly susceptible to mutagenesis. HeLa cells were infected for 90 min at 37 °C with the indicated EPEC strains, and co-precipitation analyses were carried out, as in Fig. 1. Representative gels (out of 3 independent experiments) are shown.

Next, we explored whether conservative mutations of E37 and K41 also affected Rab binding. For this purpose, EPEC-Δ*espH* complemented with EspH_E37D_, or EspH_K41R_ encoding plasmids were generated (**Fig. S1**). The mutant effector translocation into HeLa cells was confirmed (**Fig. S4**). Interestingly, these EspH mutants also failed to co-precipitate Rab8a (**Fig. 4D**). Conversely, mutating a remotely located K106 to arginine (K106R; **Fig. S1**) preserved EspH’s ability to co-precipitate the Rabs (**Fig. 4A-C**). Consistent with the co-precipitation data, colocalization analysis of translocated EspH*_K106R_*, EspH*_E37A,_* and EspH*_K41A_* with GFP-Rab8a, eGFP-Rab10, or eGFP-Rab3a, expressed in HeLa cells showed that while EspH*_K106R_* colocalized significantly with the Rabs, the other two mutants did not (**Fig. S6A-D**). These results support the understanding that residues E37 and K41 of EspH are critical for mediating the EspH-Rab interactions.

### EspH-Rab interactions are functionally relevant

Next, we investigated the impact of EspH-Rab interactions on the infected cells, focusing on previously suggested roles for these Rabs. For example, Rab8a and Rab10 have been implicated in regulating class I <u>p</u>hosphoinositide <u>3</u>-<u>k</u>inase (PI3K)/<u>m</u>ammalian target <u>o</u>f <u>r</u>apamycin <u>c</u>omplex <u>1</u> and <u>2</u> (mTORC1/C2) signaling (Lan et al, 2019; Li et al, 2010; Luo et al, 2014; Sun et al, 2010; Walia et al, 2019; Wall et al, 2017; Wang et al, 2021; Weichhart et al, 2015; Wu et al, 2023). Therefore, EspH-Rab8a interactions may affect the PI3K/Akt/mTOR signaling pathways. To address this hypothesis, HeLa cells were infected with EPEC-Δ*espH*, EPEC-Δ*espH*/pEspH*_wt_*, EPEC-Δ*espH*/pHespH*_K106R_*, EPEC-Δ*espH*/pEspH*_E37A_* and EPEC-Δ*espH*/pEspH*_K41A_*. The Akt activity level was evaluated by IB using antibodies directed against phosphorylated Ser473 of active Akt [pAkt(Ser473)]. mTORC1 activity was assessed by cell infection and immunoblotting using anti-p4EBP1 (Thr37/46) antibodies. Cell infection with EPEC-Δ*espH*/pEspH*_wt_* or EPEC-Δ*espH*/EspH*_K106R_* resulted in a significant diminishment of Akt-pSer473 and p4EBP1 (Thr37/46) levels compared to EPEC-Δ*espH* infected cells. In contrast, the levels of pAkt (Ser473) and p4EBP1 (Thr37/46) in cells infected with EPEC-Δ*espH*/pEspH*_E37A_*, or EPEC-Δ*espH*/pEspH*_K41A_* were comparable to those detected in EPEC-Δ*espH* infected cells (**Fig. 5A**). Similar results were obtained in Caco-2_BBe_ infected cells (**Fig. S7**). These results indicate that EspH inhibits the pI3K/Akt/mTORC1 signaling pathway, possibly through inhibiting Rab8a and Rab10.

**Fig. 5:**
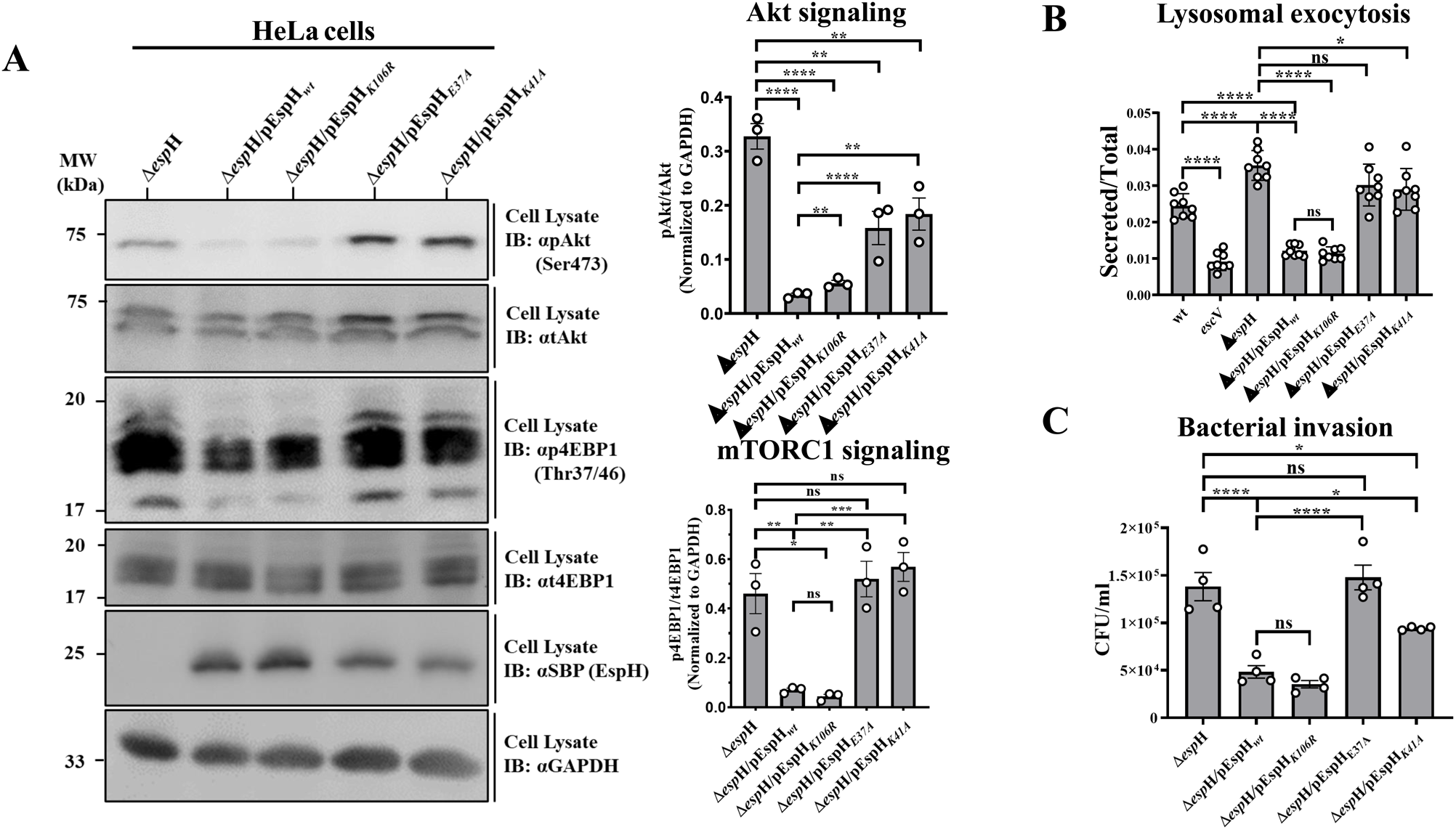
The Rab binding residues in EspH are critical for exerting Rab-related functions. HeLa cells were infected with the indicated EPEC strains for 90 min at 37 °C, and the effects on Akt and mTORC1 signaling (A), lysosome exocytosis (B), and bacterial invasion (C) were measured, as described in Materials and Methods. All experiments were repeated at least three times. Representative gels (out of at 3 independent experiments) are shown. Results are mean ±SE.

Rab3a and Rab10 have been implicated in regulating lysosomal exocytosis (Encarnacao et al, 2016; Escrevente et al, 2021; Vieira, 2018). We have recently shown that EPEC infection stimulates lysosomal exocytosis in a T3SS-dependent manner, involving the death-promoting effectors Tir, EspF, and Map (Shtuhin-Rahav et al, 2023). The observation that EspH interacts with Rab3a and Rab10 (**Fig. 1, 3, 4, S3A, and S3B**) prompted the hypothesis that the effector modulates lysosomal exocytosis. To test this, cells were infected with wild-type EPEC (EPEC-*wt*), or with an EPEC mutant deficient in T3SS activity (EPEC-*escV*), or EPEC-Δ*espH* mutant complemented or not with plasmids expressing EspH*_wt_* (EPEC-Δ*espH*/pEspH*_wt_*), or mutated EspH, including EspH*_K106R_* (EPEC-Δ*espH*/pEspH*_K106R_*), EspH*_E37A_* (EPEC-Δ*espH*/pEspH*_E37A_*), or EspH*_K106R_* (EPEC-Δ*espH*/pEspH*_K106R_*). The β-hexosaminidase activity (release) assay was applied to measure lysosomal exocytosis (Shtuhin-Rahav et al, 2023). Infection with EPEC-*escV* showed reduced lysosomal exocytosis compared to EPEC-*wt*-infected cells, confirming the involvement of the T3SS in the process. EPEC-Δ*espH* infection resulted in slightly higher lysosomal exocytosis levels than in EPEC-*wt* infected cells. In contrast, infection with EPEC-Δ*espH*/pEspH*_wt_* or EPEC-Δ*espH*/pEspH*_K106R_* caused a significant reduction in these levels (**Fig. 5B**), suggesting that injected EspH inhibits lysosomal exocytosis. In contrast, infection with EPEC strains expressing the Rab-interaction deficient mutants (EspH*_E37A_* and EspH*_K41A_*) did not affect lysosomal exocytosis (**Fig. 5B**). These data suggest that translocated EspH inhibits lysosomal exocytosis, possibly through inhibiting Rab3a or Rab10.

Studies on *Salmonella* showed that inhibiting Rab10 by the SopD effector promoted Dynamin-2 recruitment and plasma membrane scission during bacterial invasion (Boddy et al, 2021). Studies have also shown that EspH inhibits EPEC invasion by inhibiting CDC42 and Rac1 GTPases (Dong et al, 2010b; Ramachandran et al, 2022). However, these observations do not exclude the existence of other mechanisms, such as the interactions of EspH with host Rabs. Infection with EPEC-Δ*espH*/pEspH*_wt_*, or EPEC-Δ*espH*/pEspH*_K106R_* displayed significantly reduced invasion levels compared to EPEC-Δ*espH*. In contrast, infection with EPEC-Δ*espH*/pEspH*_E37A_* or EPEC-Δ*espH*/pEspH*_K41A_* did not have an effect (**Fig. 5C**). These data support the hypothesis that EspH inhibits bacterial invasion by modulating the activity of Rabs, possibly Rab10.

### Effects of EspH on Rab8a deficient cells

Given the vital role of Rab8a in PI3K/Akt/mTORC1 signaling as a mechanism that limits innate immune responses (Luo et al, 2014; Wall et al, 2017), we studied whether the EspH-mediated inhibition of these signaling pathways (**Fig. 5A**) depends on Rab8a. For this purpose, a lentiviral vector-based CRISPR/Cas9 genome editing system was used to generate HeLa cell lines deficient in Rab8a expression [Rab8a-knock-out (KO1) cells, **Fig. S8**]. HeLa cells transduced with an empty lentiviral vector and, therefore, not hampered Rab8a expression were used as controls in these experiments (Control-KO cells). To test the effects of translocated EspH, data obtained in EPEC-Δ*espH*/pEspH infected cells were compared to those of EPEC-Δ*espH* infected cells. While a significant reduction in the pAkt and p4EBP1 levels was observed in the Control-KO cells infected with EPEC-Δ*espH*/pEspH, a minor insignificant reduction was seen in the Rab8a-KO1 cells (**Fig. 6A**). These results agree with the view that EspH-Rab8a interactions play a role in the inhibition of PI3K/Akt/mTORC1 signaling. In addition, our results show that the inhibition of lysosomal exocytosis is significantly reduced in the Control-KO but not in the Rab8a-KO1 EPEC-Δ*espH*/pEspH infected cells (**Fig. 6B**), suggesting that EspH-Rab8a interactions could play a vital role in lysosomal exocytosis inhibition. However, the inhibition of bacterial invasion by translocated EspH was reduced in the Control-KO and Rab8a-KO1 cells (**Fig. 6C**), indicating that EspH-Rab8a interactions are not involved in bacterial invasion. Co-precipitation followed by IB analysis showed that eGFP-Rab3a and eGFP-Rab10 co-precipitated with translocated EspH from the Rab8a and Control-KO cells, albeit with different efficacies. While Rab3a co-precipitated at lower levels in the Rab8a-KO1 compared to the Control-KO cells, the levels of Rab10 co-precipitation were similar in the two cell lines (**Fig. 6D**). These results suggest that the interactions between EspH and Rab8a could play a role in maintaining efficient interactions with other Rabs, e.g., Rab3a.

**Fig. 6:**
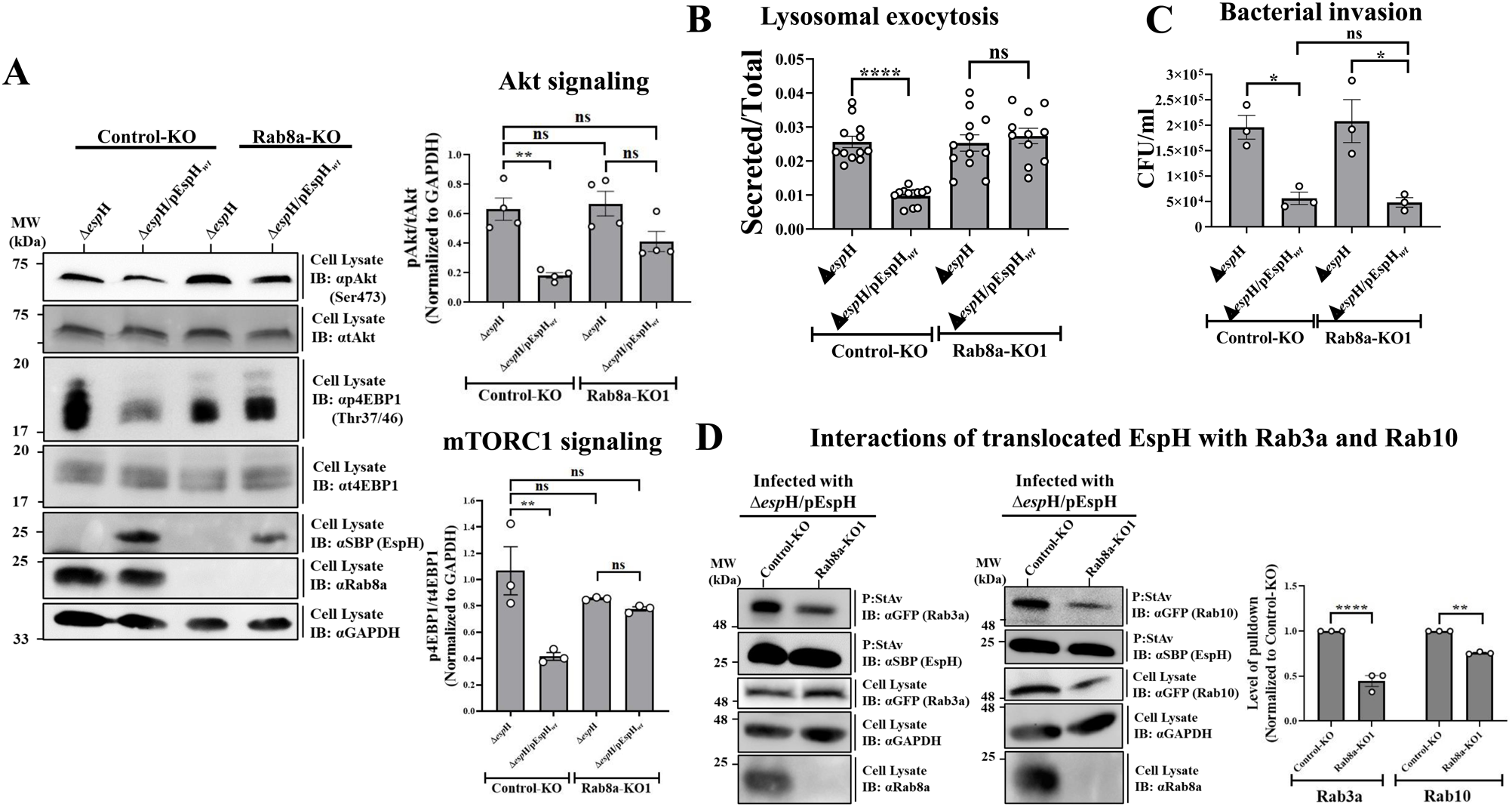
Interactions of EspH with Rab8a are essential for eliciting the AkT/mTORC1 signaling, lysosomal exocytosis, and the interactions with Rab3a. Control-KO and Rab8a-KO1 HeLa cells (**Fig. S8**) were infected with the indicated EPEC strains for 90 min at 37 °C, and the effects on Akt and mTORC1 signaling (**A**), lysosome exocytosis (**B**) and bacterial invasion (**C**), were measured, as described in Materials and Methods and Fig. 5. (**D**) Interactions of translocated EspH with Rab3a and Rab10. Control-KO and Rab8a-KO1 HeLa cells were transfected with eGFP-Rab3a and eGFP-Rab10 encoding plasmids for 48 hrs and then infected for 90 min with the indicated EPEC strains. Co-precipitation experiments were performed, as in Fig. 1. Precipitated EspH and co-precipitated Rabs were detected with anti-SBP and anti-GFP antibodies, respectively. Anti-GAPDH antibodies were used to assess the protein load. The level of the co-precipitated Rabs was calculated by measuring the intensity of the co-precipitated protein band normalized to the intensity of the protein band detected in the cell lysate and the intensity of the GAPDH band. The values obtained were further normalized to the Control-KO levels. All experiments were repeated at least three times. Representative gels (out of at 3 independent experiments) are shown. Results are mean ±SE.

### EspH-Rab interactions do not play a role in stimulating host cytotoxicity and filopodia repression

Studies have shown that translocated EspH induces host cell cytotoxicity and inhibits transient filopodia formation, functions attributed to Rho GTPase inhibition (Ramachandran et al, 2022). Infection of HeLa cells with EPEC-Δ*espH*/pEspH*_E37A_* or EPEC-Δ*espH*/pEspH*_K41A_* did not affect host cell cytotoxicity (**Fig. 7A**) or filopodia repression (**Fig. 7B**) compared to EspH*_wt_* infected cells, suggesting that Rab binding by EspH is not involved in these processes. These results further signify the existence of two distinct functional motifs in EspH: one is the C-terminal 38aa segment that binds ABR to downregulate Rho GTPases, inducing cell cytotoxicity and repressing filopodia formation (Ramachandran et al, 2022), and the other is the Rab binding motif located upstream of this segment, involved in modulating Rab GTPases and their role in Akt/mTORC1 signaling, lysosomal exocytosis and bacterial invasion.

**Fig. 7:**
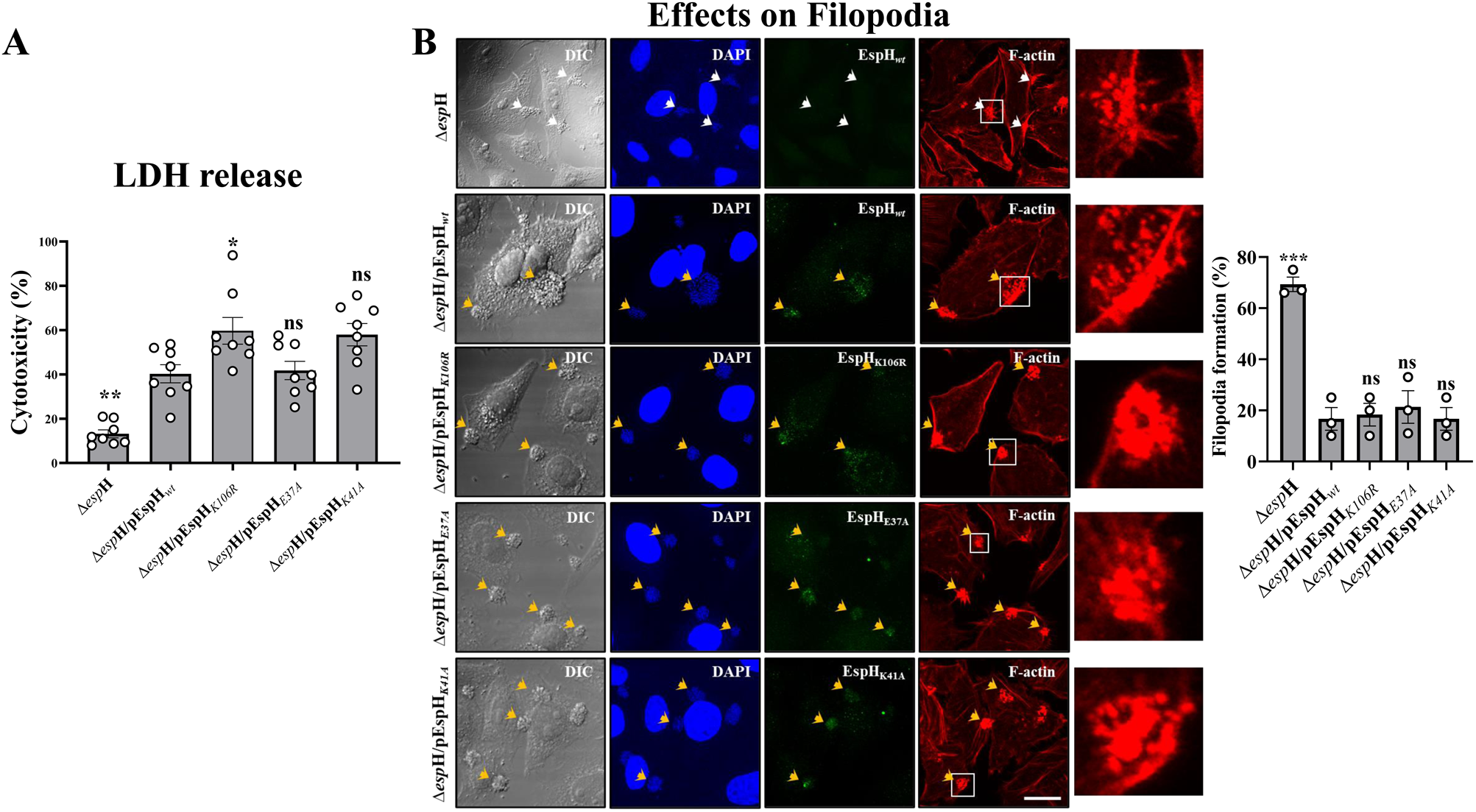
The effects of mutations in EspH used to study the interactions with Rab GTPases on host cell cytotoxicity and filopodia formation. HeLa cells were infected for 90 or 15 min with the indicated EPEC strains, and the impact on cell cytotoxicity using the LDH release assay **(A)** or filopodia formation by confocal cell imaging **(B)**, respectively, was measured as described in Materials and Methods. For cell imaging, EspH was immunostained by anti-SBP antibodies, host cell nuclei and bacterial microcolonies were visualized by DAPI staining, and F-actin was observed by Phalloidin Texas Red staining. Representative images from 3 independent experiments are shown. Results are mean ±SE. Statistical significance tests were performed versus the EPEC-Δ*esp*H/EspH*_wt_* infected cells.

### A PBD of EspH mediates effector localization at plasma membrane infection sites and interactions with Rab8a

A common conserved GKx<u>Y</u>x_n_F PI-binding domain (PBD) with a critical tyrosine (underlined) has been identified in type III-secreted effectors, mediating high-affinity interactions with PI-enriched membrane platforms (Salomon et al, 2013). Interestingly, we identified the conserved domain in EspH expressed by several pathogenic *E. coli* species, including EPEC, and thus mutated its conserved tyrosine at position 68 to alanine (Y68A; **Fig. S1**). The translocation of EspH*_Y68A_* into HeLa cells infected with EPEC-Δ*espH*/pEspH*_Y68A_* was confirmed (**Fig. S4**). HeLa cells expressing eGFP-PH-Akt [a reporter of PI(3,4)P_2_ and PI(3,4,5)P_3_] or eGFP-PH-TAPP1 [a reporter of PI(3,4)P_2_] (Posor et al, 2022) were infected with EPEC-Δ*espH*/pEspH*_wt_* or EPEC-Δ*espH*/pEspH*_Y68A_*. Confocal imaging showed that translocated EspH colocalized extensively with each PI reporter in large patches at the plasma membrane infection sites. The translocated EspH*_Y68A_* neither clustered nor colocalized with the PI sensors at the infection sites (**Fig. 8A**). These results may agree with previous reports, suggesting the existence of PI-enriched membrane platforms at infection sites (Campellone, 2010; Sason et al, 2009; Smith et al, 2010) with which EspH interacts via its PBD. These interactions may promote EspH preferential localization and PI clustering at these sites.

**Fig. 8:**
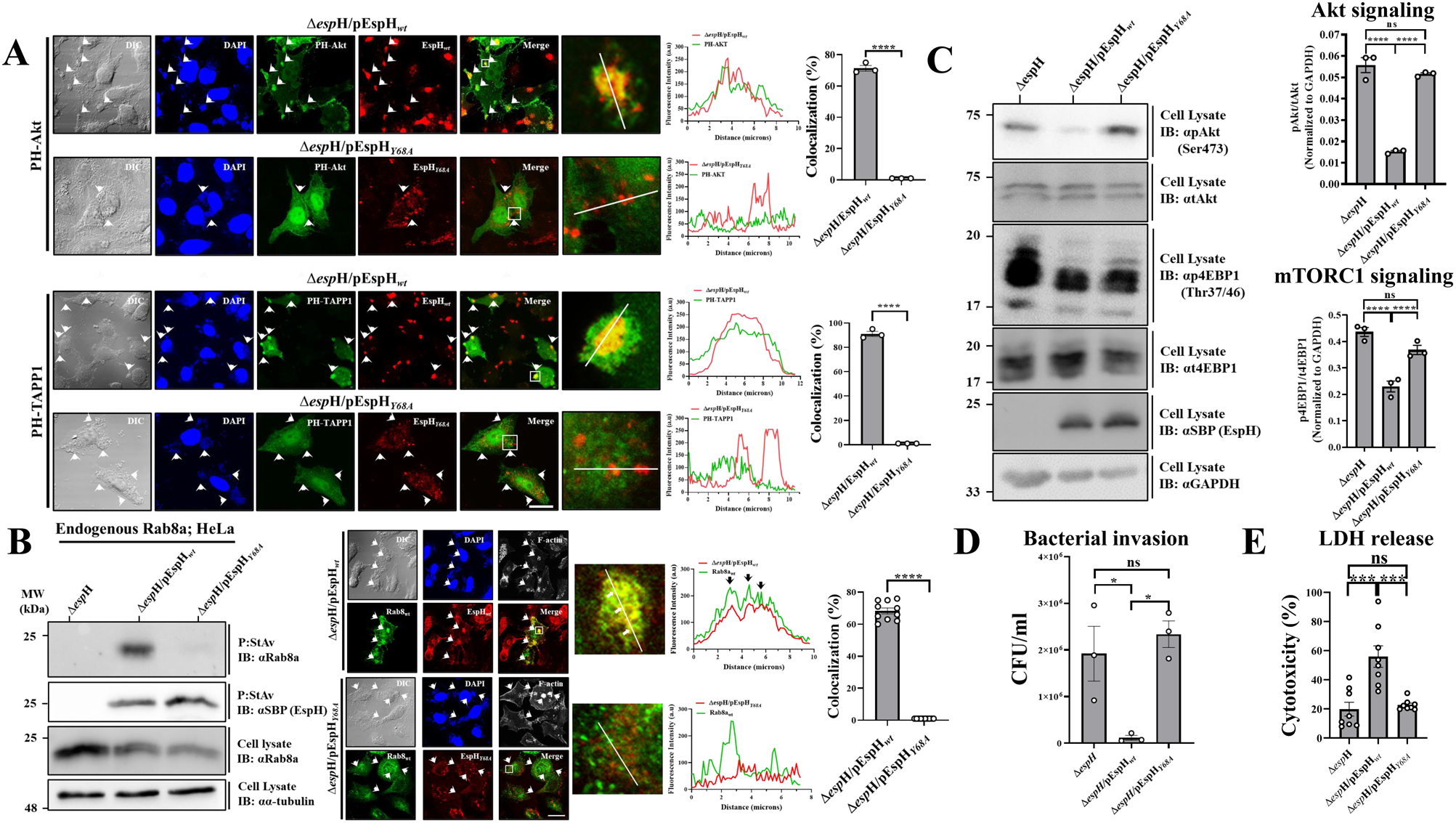
The EspH PBD is critical for EspH localization at infection sites, PI clustering, and interactions with Rab8a. **(A). PI clustering at infection sites.** HeLa cells were transfected with eGFP-PH-Akt (upper) or eGFP-PH-TAPP1 encoding plasmids (lower). Eighteen hours post-transfection, cells were infected with EPEC-Δ*espH*/pEspH*_wt_* or EPEC-Δ*espH*/pEspH*_Y68A_* strains for 30 min at 37 °C. Cells were fixed, permeabilized, and immunostained with anti-SBP antibodies to visualize EspH (red). Cells were then stained with DAPI (blue) to visualize host nuclei and adhered bacterial microcolonies (indicated with arrowheads) and analyzed by confocal microscopy. Colocalization analysis was performed as described in Fig. 2. (B) Interactions with Rab8a; *Analysis by co-precipitation (left)*. HeLa cells were infected for 90 min with the indicated EPEC strains, and co-precipitation experiments detecting the endogenous Rab8a were performed, as in Fig. 1. **Analysis by colocalization (right).** HeLa cells were transfected with a GFP-Rab8a*_wt_* encoding plasmid and infected with the indicated EspH-expressing EPEC strains. Colocalization analyses were performed, as in Fig. 2. **(C) Effects on Akt/mTORC1 activities.** HeLa cells were infected for 90 min with the indicated EPEC strains, and the assay was performed as in Fig. 5**. (D) Effects on bacterial invasion.** HeLa cells were infected for 90 min at 37 °C with the indicated EPEC strains, and the invasion assay was performed as described in Materials and Methods. (**E**) **Effects on host cytotoxicity.** HeLa cells were infected with the indicated strains for 90 min at 37 °C, and the LDH release assay was used to evaluate the impact on cell cytotoxicity. All experiments were repeated at least three times. Representative gels are shown. Results are mean ±SE.

Next, we examined whether the PBD is essential for the EspH-Rab8a interactions. HeLa cells infected with EPEC-Δ*espH* (a negative control) EPEC-Δ*espH*/pEspH*_wt_* or EPEC-Δ*espH*/pEspH*_Y68A_*, were subjected to the Rab co-precipitation method. The endogenous Rab8a co-precipitated with EspH*_wt_* but not EspH*_Y68A_* (**Fig. 8B, left**). Confocal imaging showed significant colocalization between GFP-Rab8a and translocated EspH*_wt_*, but not EspH*_Y68A_*, in large clusters at infection sites (**Fig. 8B, right**). Notably, the Alphafold predicted structure of EspH-Rab8a suggested that Y68 of the EspH PBD is distinct from the Rab binding interface (**Fig. S9**). While infection with EPEC-Δ*espH*/pEspH*_wt_* showed a significant reduction of pAkt and p4EBP1 levels compared to EPEC-Δ*espH*, cells infected with EPEC-Δ*espH*/pEspH*_Y68A_* displayed an impaired ability to dephosphorylate Akt and 4EBP1 (**Fig. 8C**). While infection with EPEC-Δ*espH*/pEspH*_wt_* caused significant inhibition of bacterial invasion compared to EPEC-Δ*espH* infected cells, infection with EPEC-Δ*espH*/pEspH*_Y68A_* did not yield an effect (**Fig. 8D**). Unlike translocated EspH*_wt_* which, as previously demonstrated (Ramachandran et al, 2022; Tu et al, 2003), induced cytotoxicity, translocated EspH*_Y68A_* had no impact on host cytotoxicity (**Fig. 8E**). These data argue that the PBD mediates the interactions of EspH with host Rab8a. They also indicate that the interactions are vital for exerting EspH-dependent suppression of Akt, mTORC1, bacterial invasion into the host cells, and induction of host cytotoxicity. Thus, the interactions of EspH with plasma membrane PIs are vital for exerting all its reported functions.

## Discussion

EspH, an effector protein known for its ability to disrupt the host cell actin cytoskeleton by inhibiting Rho GTPases (Dong et al, 2010b; Ramachandran et al, 2022; Ramachandran et al, 2018; Roxas et al, 2018; Tu et al, 2003; Wong et al, 2012a; Wong et al, 2012c), is shown here to interact with several Rab GTPases, including Rab8a, Rab10, Rab3a, and Rab12 (**Figs. 1-4**). In this regard, EspH joins a list of bacterial effector proteins shown to bind multiple host Rabs (Boddy et al, 2021; Cheng et al, 2012; Gan et al, 2020; Goody et al, 2012; Lian et al, 2021; Mousnier et al, 2014; Schoebel et al, 2011; Spano et al, 2016). The ability of EspH to bind these Rabs and precipitate them from cell lysates (**Fig. 2 and Figs. S2A and S2B**) suggests that the binding is firm, an idea consistent with reports for other effector-Rab interactions (Dubreuil & Segev, 2011; Schoebel et al, 2011). The mechanism by which EspH affects the host Rabs is not clear. However, assuming that it inhibits normal signaling pathways by binding the active Rab forms (**Fig. 2 and Fig. S3A and S3B**) would be conceivable. It would also be reasonable to hypothesize that EspH-mediated inhibition of Akt/mTORC1 signaling by binding active Rab8a (**Fig. 6A**) causes the inhibition of downstream cytokine secretion and inflammatory responses, as described earlier in the case of LPS-treated macrophages (Luo et al, 2014; Wall et al, 2017) and for the *Salmonella* SopD effector protein (Lian et al, 2021).

We have recently shown that lysosomal exocytosis is linked to the translocation of death-promoting effectors, e.g., Tir, EspF, and Map (Shtuhin-Rahav et al, 2023). Here, we show that translocated EspH inhibited lysosomal exocytosis in a Rab8a-dependent fashion (**Fig. 6B**). mTORC1 signaling has been tightly linked to lysosomal positioning and trafficking (Hong et al, 2017; Mutvei et al, 2020; Puertollano, 2014). Hence, downregulation of mTORC1 activity by EspH-Rab8a interactions (**Fig. 6A**) may play a role in lysosomal exocytosis inhibition (**Fig. 6B**). The efficiency of Rab3a to co-precipitate with EspH is reduced in the absence of Rab8a (**Fig. 6D**), suggesting that the capacity of the effector to interact with Rab3a depends on Rab8a binding. As Rab3a has also been proposed to modulate lysosomal exocytosis (Encarnacao et al, 2016; Vieira, 2018), the reduced capacity of EspH to inhibit lysosomal exocytosis in the Rab8a-deficient cells may be contributed by reduced Rab3a binding. EspH induces cell cytotoxicity, imposing cell rounding and plasma membrane damage (Ramachandran et al, 2022). Lysosomal exocytosis regulated by Rab10 and Rab3a has been implicated in membrane repair (Vieira, 2018). The blockade of host membrane repair by EspH may represent a novel type of cell stress imposed by the effector.

AlphaFold identified residues in an α-helix of EspH (E37 and K41) that bind the K and D residues in the β-strand interswitch region of all the indicated Rabs, suggesting this is a common characteristic in Rab binding (**Fig. 3**). Crystallographic studies have found similar effector-Rab interactions: e.g., α-helices in *Legionella* LidA that bind the two switches and the interswitch of Rab1 via polar aa and salt bridges (Cheng et al, 2012). The structure of the complex (PDBid 3SFV) shows interactions between K49 and D47 on a β-strand of Rab1 to the corresponding D235 and K239 on LidA, similar to the E37 to K and K41 to D of EspH-Rab interactions. Similarly, the *Salmonella* SopD effector displays a large interaction interface with Rab8a, including experimentally identified E293 and K285 in an α-helix in the effector protein that interacts with K58 and Q60, respectively, in the interswitch β-strand of Rab8a (Lian et al, 2021).

A hydrophobic interaction was seen between the hydrophobic triad (F45, Y77, and W62) in human Rab8a and LidA (Schoebel et al, 2011). We also found aromatic interactions between F46 and W63 of the hydrophobic triad of Rab10 and F59 and W76 of Rab3a, with F10 of EspH (**Fig. 3A and 3B**). These interactions may contribute to stronger binding, which may be why the PAE confidence is higher for these complexes in the AlphaFold structures (**Fig. 3C**). In conclusion, the Rab-effector structural interface, exemplified in the case of EspH-Rab binding and found in other bacterial effector-Rab interfaces, may reflect a common mode of binding that enables the functional manipulation of multiple Rabs by these effectors and contributes to pathogenesis.

Our data indicate that plasma membrane PIs play a role in enabling the EspH-Rab interactions (**Fig. 8**). PI sensors and translocated EspH*_wt_* clustered significantly at the infection sites. The clustering effect was significantly reduced in the presence of EspH*_Y68A_* (**Fig. 8A**). Interestingly, Tir has been suggested to build-up of PI(3,4)P_2_ and PI(3,4,5)P_3_ enriched platforms at plasma membrane infection sites (Campellone, 2010; Sason et al, 2009; Smith et al, 2010). EspH may further augment this effect by binding and clustering host PIs via its PBD. Plasma membrane PI domains have been implicated in the recruitment and regulation of Rab and Rho GTPases spatiotemporal signaling (Croise et al, 2014; Jean & Kiger, 2012; Viaud et al, 2012). Therefore, the EspH-PI interactions at bacterial infection sites may represent another mechanism by which the effector protein impacts the structure and function of host Rab and Rho GTPases.

As an extracellular bacterial pathogen, EPEC elicits mechanisms that block its invasion (phagocytosis) into the host cells, enabling it to remain extracellular (Goosney et al, 1999). These mechanisms involve the inhibition of Rho GTPases by EspH (Dong et al, 2010b; Ramachandran et al, 2022), EspG-mediated counteracting of the WAVE regulatory complex (Humphreys et al, 2016), and inhibition of PI3K-dependent signaling pathways (Celli et al, 2001). Here, we propose another anti-phagocytic mechanism involving EspH binding to host Rabs (**Fig 5C**).

Our studies suggest that the effector protein EspH of A/E pathogens contains PI, ABR, and Rab GTPase binding domains. EspH utilizes a PBD to confine its localization at plasma membrane infection sites. These interactions facilitate the binding and downregulation of the activity of host Rho (through binding ABR) and Rab GTPases to disrupt the actin cytoskeleton, lysosomal trafficking, and immune signaling pathways, thereby contributing to bacterial pathogenesis (Ritchie & Waldor, 2005).

## Materials and Methods

### Bacterial strains, antibodies, plasmids, primers

Bacterial strains, antibodies, plasmids, primers, and reagents used in this study are listed in the **supplementary Table S1**.

### Cell culture and transfection of cells with plasmid DNA

HeLa and CaCo-2_BBe_ cells (semi-polarized) (see **Table S1**) were cultured, as previously described (Ramachandran et al, 2018). Plasmids were transiently transfected into HeLa cells (∼ 60% confluence) for 48 hrs unless otherwise indicated using the TransIT-X2 Transfection Reagent protocol.

### Construction of EspH mutants

The mutations were constructed on the pSA10-EspH*_wt_*-6xHis-SBP encoding plasmid using PCR amplification of the vector and inserts and ligation using Gibson assembly. The vector was linearized using the 1F and 1R oligonucleotides. The oligonucleotides 2F and 2R or 3F and 3R were used to mutate glutamic acid at position 37 to alanine or aspartic acid and generate the pSA10-EspH*_E37A_* and pSA10-EspH*_E37D_* encoding plasmids, respectively. Oligonucleotides 4F and 4R or 5F and 5R were used to mutate lysine at position 41 to alanine or arginine and generate the pSA10-EspH*_K41A_* or pSA10-EspH*_K41R_* encoding plasmids, respectively. Oligonucleotide 6F and 6R were used to mutate lysine at position 106 to arginine and generate the pSA10-EspH*_K106R_* plasmid. Oligonucleotides 7F and 7R were used to mutate tyrosine at position 68 to alanine and create the pSA10-EspH*_Y68A_* plasmid. Nucleotide sequences of all constructs were confirmed by the Genomic Technologies Facility (https://www.bio.huji.ac.il/en/units_the_national_center_for_genomic_technologies) using SANGER sequencing. All the EspH mutant plasmids were electroporated at 1.85 kV/25 µF/200 Ohm, using the BioRad electroporator (GENE PULSER II) into the EPEC-Δ*esp*H strains to generate the plasmid complemented bacterial strains.

### Bacterial pre-activation and cell infection

Before infection, the T3SS of bacterial strains was activated in plain high glucose Dulbecco’s Modified Eagle Medium (DMEM) for 3 hrs in the CO_2_ incubator (37 °C, 5% CO_2_, 95% humidity), as described (Ramachandran et al, 2020). The expression of EspH in EPEC-Δ*esp*H strains was induced by supplementing the activation medium with isopropyl-β-D-thiogalactopyranoside (IPTG; 0.05 mM for inducing EspH*_wt_* expression, 0.1mM for inducing the EspH mutant expression, and as indicated in the Figures) during the last 30 min of activation. Cell infection was performed with the pre-activated infection medium in the CO_2_ incubator at 37 °C for the indicated times. Bacterial infection was performed at a multiplicity of infection (MOI) of ∼100.

### SDS-PAGE and Immunoblotting

SDS-PAGE and IB were performed as described (Shtuhin-Rahav et al, 2023). Band intensity was measured using Fiji (NIH).

### Effector translocation assay

The effector translocation assay was performed as described (Ramachandran et al, 2020). Approximately 2×10^5^ HeLa cells/well were seeded in a 6-well plate and cultured for 48 hrs until reaching ∼ 70% confluence. Cells were then infected for 90 min at 37 °C with the indicated EPEC strains, and IPTG was used to induce EspH expression. Following infection, cells were washed three times with ice-cold PBS and lysed in 60 µl of ice-cold lysis buffer [100 mM NaCl, 1 mM EDTA, 10 mM Tris-HCl, pH 7.4, 0.5% (vol/vol) NP-40] supplemented with protease and phosphatase inhibitors. Following 3 min incubation on ice, the lysate was pipetted up and down and then centrifuged (10000 g, 4 °C, 10 min). The supernatant (detergent-soluble) and pellet (detergent-insoluble) fractions were isolated. The pellet was resuspended in 60 µl of the lysis buffer. Twenty microliters of 4x SDS-PAGE sample buffer were added to each lysis buffer containing a fraction. The samples were then heated (95 °C; 10 min) and analyzed by SDS-PAGE, followed by IB. The presence of EspH in the fractions was detected by anti-SBP antibodies. Anti-α-tubulin antibodies were used to assess the lysate protein load.

### Analyzing EspH-Rab interactions by co-precipitation (pulldown) assays

HeLa or Caco-2_BBe_ cells cultured on 15-cm plates (70 % confluence) were infected for 90 minutes with pre-activated EPEC strains. Cells were washed three times with ice-cold PBS and lysed in ice-cold lysis buffer [50mM Tris (pH 7.4), 150 mM NaCl, 0.5% NP-40] supplemented with protease and phosphatase inhibitors. Lysates were centrifuged (5,000 g, 15 min, 4 °C), and the Bradford reagent was used to determine the supernatants’ protein concentration. EspH was precipitated (P) from an equal amount (∼5mg) of cell lysates by incubation with 60 µl of Streptavidin (StAv) agarose beads (50% slurry pre-washed with lysis buffer) for 3 hrs at 4°C with end-to-end rotation. Beads were washed three times with lysis buffer by centrifugation (300 g, 2 min, 4 °C), dried using Hamilton’s syringe, and subjected to SDS-PAGE followed by IB. Pulled-down EspH and co-precipitated Rabs were detected using anti-SBP and appropriate anti-Rab or epitope-tagged antibodies.

### Immunofluorescence labeling of permeabilized cells

The procedures applied were performed as described (Haritan et al, 2023; Kassa et al, 2019).

### Fluorescence microscopy and colocalization analyses

Cells were processed and imaged by an Olympus FV-1200 laser scanning confocal microscope equipped with a 60× oil immersion objective (NA, 1.42), as described (Haritan et al, 2023). Colocalization analyses using the intensity profile tool of Fiji (NIH) were performed, as described (Haritan et al, 2023). Scale bars = 5 μm.

### The β-hexosaminidase activity (release) assay

Lysosomal exocytosis was evaluated using the β-hexosaminidase activity measurements in HeLa-infected cells (Shtuhin-Rahav et al, 2023).

### Bacterial invasion and filopodia formation measurements

Bacterial invasion and the induction of transient filopodia in infected HeLa cells were measured, as described (Ramachandran et al, 2022).

### Akt/mTORC1 activity measurements

HeLa or Caco-2_BBe_ cells cultured on 10-cm plates (70% confluence) were infected for 90 min with pre-activated EPEC. Cells were then washed three times with ice-cold PBS and lysed in ice-cold lysis buffer [50 mM Tris (pH 7.6), 150 mM NaCl, 10% glycerol, 1% Triton X-100, 1 mM EDTA, 2 mM MgCl_2_, 100 mM NaF, 200 μM NaVO_4_, 1 mM PMSF, 1 µM leupeptin and 1 µM aprotinin]. Lysates were centrifuged (10,000g, 10 min, 4 °C), and the protein concentration of the supernatants was determined using the Bradford reagent. An equal amount of protein (∼100 µg) was analyzed by SDS-PAGE, followed by IB. Active Akt and mTORC1 levels were evaluated by probing the blots with antibodies directed to phosphor (p) Akt (Ser473) or phosphor (p) 4E-BP1, respectively. The band intensities of phosphorylated proteins were first normalized to total (t) Akt or 4E-BP1 protein band intensities, then to the glyceraldehyde-3-phosphate dehydrogenase (GAPDH) protein band.

### Establishing a HeLa Rab8a knock-out (KO) cell line by a lentivirus-based CRISPR/Cas9 genome editing system

#### Lentivirus preparation

The gRNA sequences (8F’ and 8R’; see **Table S1**) targeting the human Rab8a gene were cloned into lentiCRISPR V2 lentiviral vector (**Table S1)** as described by the Zhang laboratory (https://media.addgene.org/data/plasmids/52/52961/52961-attachment_B3xTwla0bkYD.pdf). To produce the lentiviruses, HEK293T/SF17 cells were seeded (3.8×10^6^ cells/plate) onto 10-cm plate and grown for 24 hrs in a CO_2_ incubator. Then, the cells were treated with 10 µl of 25 µM chloroquine and incubated in a CO_2_ at 37 °C incubator for 5 hrs. Thereafter, the cells were transfected with a mixture of psPAX2 (10 µg), pMD2.G (6 µg), and either gRNA-containing lentiCRISPRV2 (10 µg) or empty lentiCRISPRV2 (10 µg) plasmids, using polyehyleneimine (PEI; DNA: PEI 1:3 w/w ratio), and incubated in the CO_2_ incubator for 16 hrs. Then, the transfection medium was replaced after 6 hrs with a complete DMEM lacking antibiotics to allow viral particle production and release for 48 or 96 hrs. The virus-containing medium was harvested, pooled, and centrifuged (500 x g, 10 min, 22 °C), and the supernatant was clarified by passing through a 0.45 μm filter unit. Viral particles were concentrated by gently mixing one volume of PEG8000 solution 32% w/v in PBS with 3 volumes of clarified supernatant and incubated for 30 min at 4 °C, followed by centrifugation (1,500 x g 45 min 4 °C). The viral pellet was dissolved in 12 ml of complete DMEM without antibiotics.

#### Cell infection and clone isolation

HeLa cells were seeded in a 6-well plate (1.5×10^5^ cells/well) and incubated for 24 hrs in the CO_2_ incubator before infection. The cells were then infected with 2 ml of the concentrated lentiviral preparation supplemented with 8 ug/ml polybrene and incubated for 24 hrs in the CO_2_ incubator. The cell infection treatment was repeated for three successive days. After the third infection, cells were washed with PBS and selected with complete DMEM containing puromycin (3 µg/ml) for 48 hrs. Cells were then cultured in 10-cm plates at 50 cells/plate in DMEM containing puromycin (3 µg/ml) and allowed to grow in the CO_2_ incubator for approximately ten days until single cell colonies were identified. Individual cell colonies were then picked by trypsinization and expanded for analysis of Rab8a expression by IB (**Fig. S8**).

### Lactate dehydrogenase (LDH) cytotoxicity assay

HeLa cells (10,000 cells/well) were seeded on a 96-well plate and cultured for 48 hrs in a CO_2_ incubator until reaching ∼70% confluence. Cells were infected with EPEC, and the LDH release assay was applied to the cell culture medium using the manufacturer’s CytoTox 96® Non-Radioactive Cytotoxicity Assay. The cytotoxicity was calculated as follows: 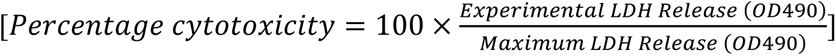, whereby ‘Experimental LDH Release’ denotes the LDH release into the medium bathing the cells; ‘Maximum LDH Release’ represents the LDH levels in cells lysed with 1X lysis solution provided by the kit.

### Statistical analysis

The GraphPad Prism v. 8.4.3 software was used for statistical analysis and graphing. A one-way ANOVA followed by Bonferroni’s multiple-comparison test was applied to determine the statistical significance. The significance is indicated by asterisks, as follows: ****p≤0.0005; ***p>0.0005; **p<0.005; *p>0.005; ns, non-significant p>0.05. A p-value < 0.05 indicates a statistically significant difference.

## Acknowledgments

RRS and IN are the Dr. Willem Been Legacy Fellowship recipients. This research was supported by the Israel Science Foundation (grant No. 1671/19).

## Data Availability Section

This study includes no data deposited in external repositories.

## Supplementary Figure legends

**Fig. S1: Clustal aa sequence alignment of EspH effector proteins expressed by pathogenic *E. coli*.** The Genebank names of EspH are indicated on the left, and amino acid (aa) numbers are shown on the right. The corresponding pathogenic *E. coli* strains are as follows: ECSP_4679-O157:H7 str TW14359 (EHEC); ECO103_3614-O103:H2 str 12009 (VTEC); ECO26_5275-O26:H11 str 11368 (STEC); ROD_29891-*Citrobacter rodentium* ICC168; ECO111_3748-O111:H-str 11128 (EHEC); E2348C_3944-O127:H6 str E2348/69 (EPEC). The CLUSTAL O (1.2.4) multiple sequence alignment program (https://doi.org/10.1093/nar/gkac240) was used. * residues indicate positions having a single and fully conserved residue. : indicates conserved displaying strongly similar properties (score > 0.5 on the PAM 250 matrix). · indicates semi-conserved residues showing weakly similar properties (score ≤ 0.5 on the PAM 250 matrix). Blanks indicate non-conserved residues. The black vertical boxes indicate the residues mutated in this study (i.e., E35A/D, K41A/R, Y68A, and K106R). The yellow highlighted region indicates the common GKxYx_n_F PI-binding domain (PBD). The horizontal box indicates the C-terminal 38 aa segment ABR binding motif.

**Fig. S2: Rab8a is co-precipitated with translocated EspH*_Δ130-168_*.** Experiments were performed as described in **Fig. 1** and Materials and Methods. A representative gel (out of 3 independent experiments) is shown.

**Fig. S3: Translocated EspH*_wt_* interacts with active Rab3a (A) and Rab10 (B).** Experiments were performed as described in Materials and Methods and **Fig. 2**. Rab3a_T36N_, Rab3a_Q81L_, Rab10_T23N_, and Rab10_Q68L_ represent the dominant negative (GDP-bound) and constitutively active (GTP-bound) Rab forms, respectively. Representative gels and confocal images (out of 3 independent experiments) are shown.

**Fig. S4: Translocation of EspH mutants.** HeLa cells were infected for 90 min at 37 °C with indicated EPEC strains. The different concentrations of IPTG were used to induce effector protein expression levels. The effector translocation assay was applied, as described in Materials and Methods. EspH was detected using anti-SBP antibodies. Anti-α-tubulin (αα-tubulin) antibodies were used to evaluate protein loading.

**Fig. S5: The Rab binding residues E37 and K41 of EspH are critical for binding endogenous Rab8a in Caco-2_BBe_ cells.** Caco-2_BBe_ cells were infected with indicated EPEC strains for 90 min at 37 °C, and a co-precipitation experiment was carried out, as described in Materials and Methods and **Fig. 4**.

**Fig. S6: The Rab binding residues in EspH are critical for recruiting Rabs to infection sites.** HeLa cells expressing GFP-Rab8a*_wt_* (**A**), eGFP-Rab10*_wt_* (**B**), or eGFP-Rab3a*_wt_* (**C**) were infected with the indicated EPEC strains for 90 min at 37 °C. Cells were subjected to EspH immunostaining using anti-SBP antibodies, DAPI (to visualize host nuclei and bacterial microcolonies (pointed with arrowheads), Phalloidin CF-647 (to visualize F-actin), and confocal imaging followed by colocalization analysis (**D**). Results are mean ±SE.

**Fig. S7: The Rab binding residues in EspH are critical for exerting Akt/mTORC1 signaling in Caco-2_BBe_ cells.** Caco-2_BBe_ cells were infected with indicated EPEC strains for 90 min at 37 °C, and the effects on Akt and mTORC1 signaling were assessed, as described in **Fig. 5** and Materials and Methods. Results are mean ±SE.

**Fig. S8: Analysis of CRISPR/Cas9 Rab8a KO cells by IB.** The Control-KO and 4 single Rab8a-KO clone cells were lysed and analyzed by IB for the presence of endogenous Rab8a using anti-Rab8a antibodies. Anti-GAPDH antibodies were used to evaluate protein loading. Rab8a-KO1 was used for further experiments.

**Fig. S9: AlphaFold predicted structure of EspH-Rab8a shows that the Y68 residue of the EspH PBD is distinct from the Rab binding site.** The complex of EspH-Rab8a was modeled using AlphaFold-Multimer-v2.0 (see also **Fig. 3**). The EspH (pink) and the Rab GTPase (gray) structures are depicted. The switch I, interswitch, and switch II Rab domains are shown in maroon, dark green, and navy blue, respectively. The interface area shows the predicted interacting E37 and K41 of EspH (yellow) and K46 and D44 of Rab8a interswitch region (green). The PBD of EspH (orange) is shown with Y68 (light green).

---

**Supplementary Table S1.**
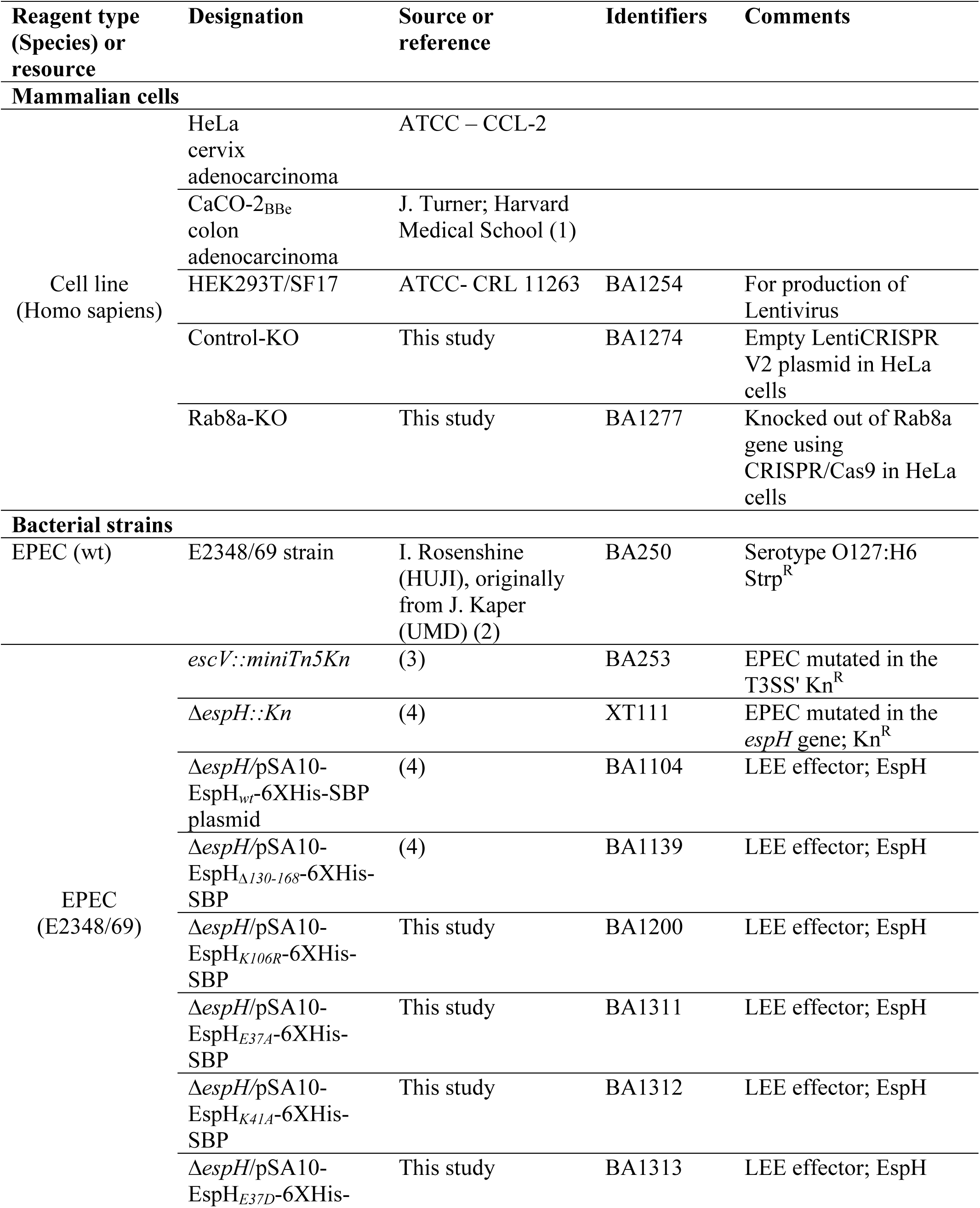
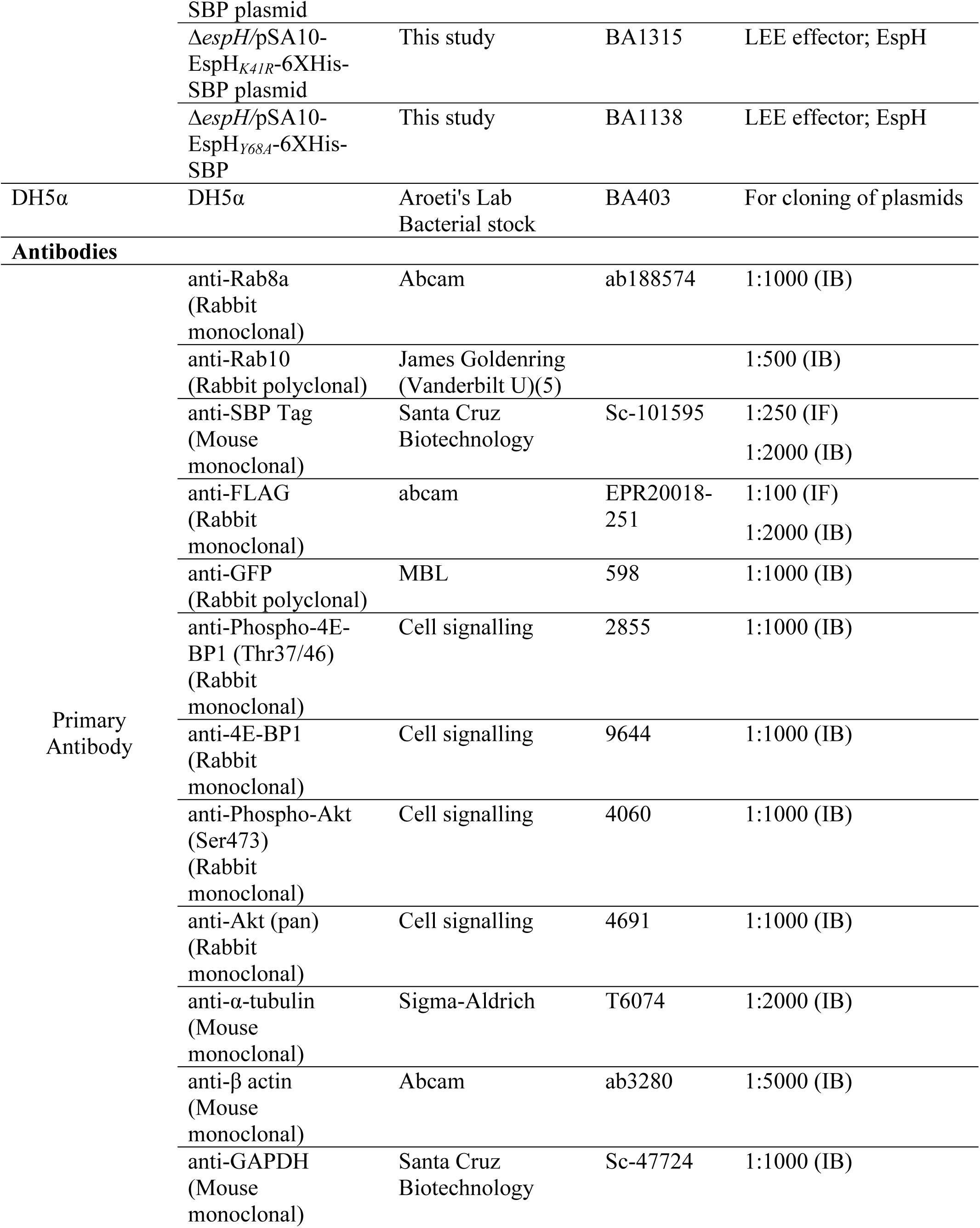
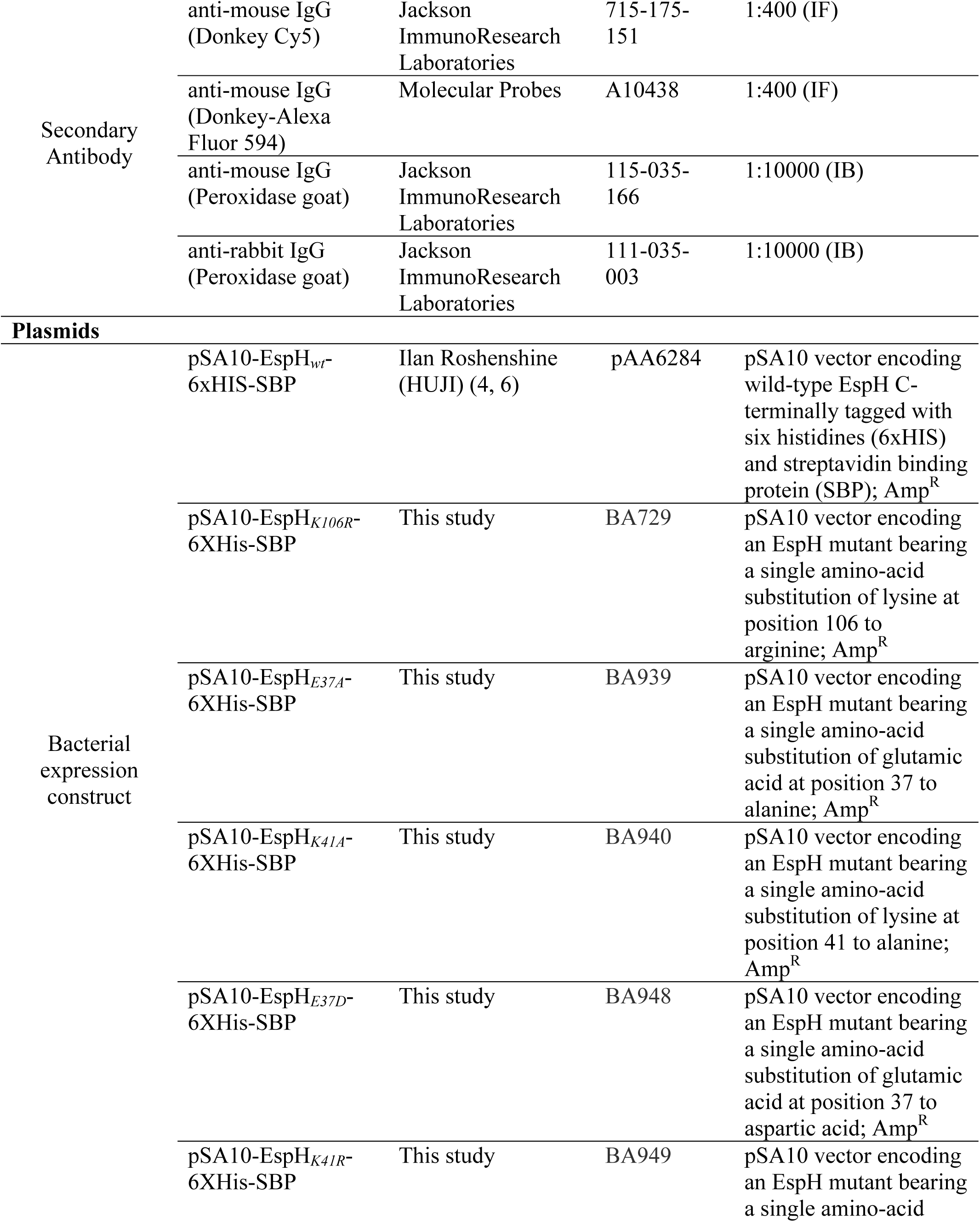
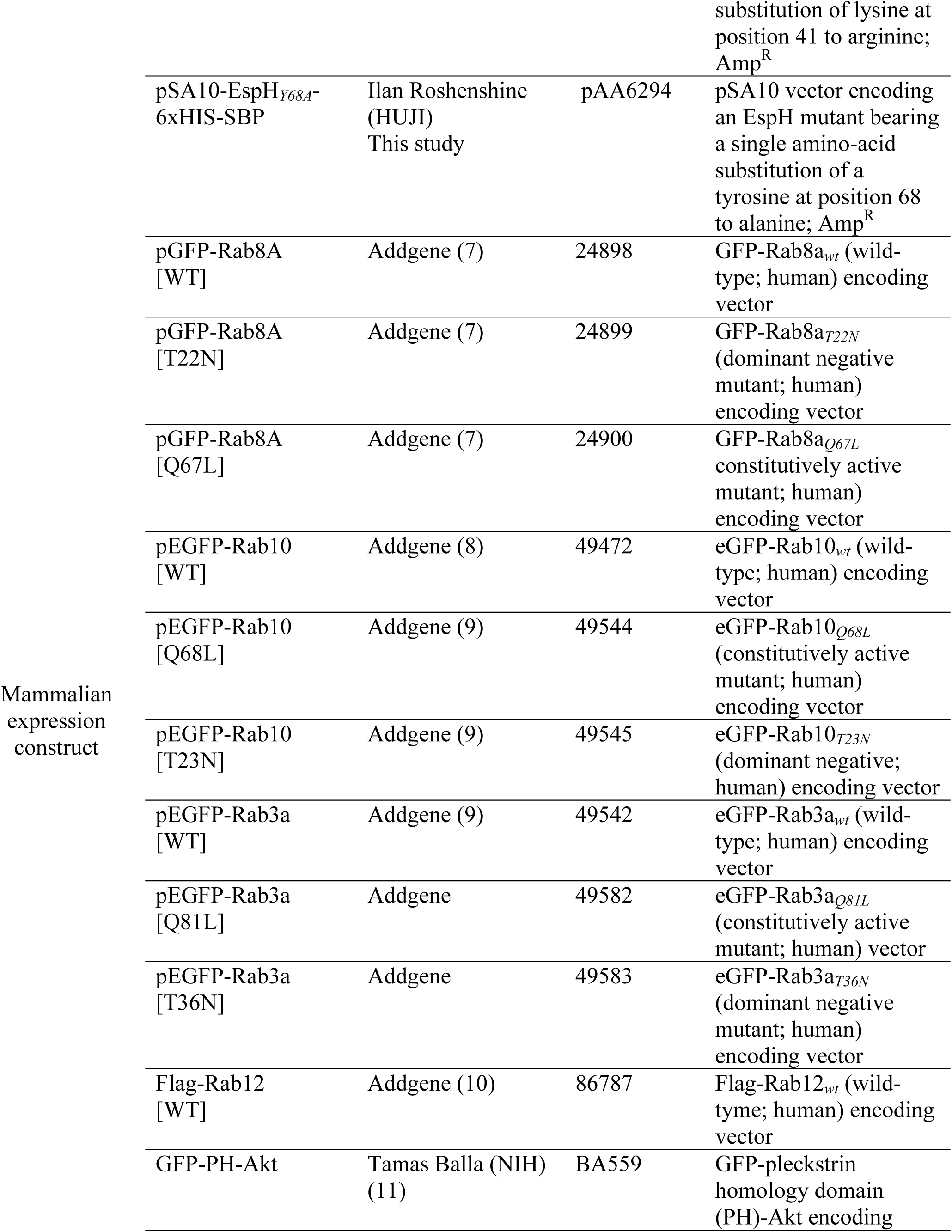
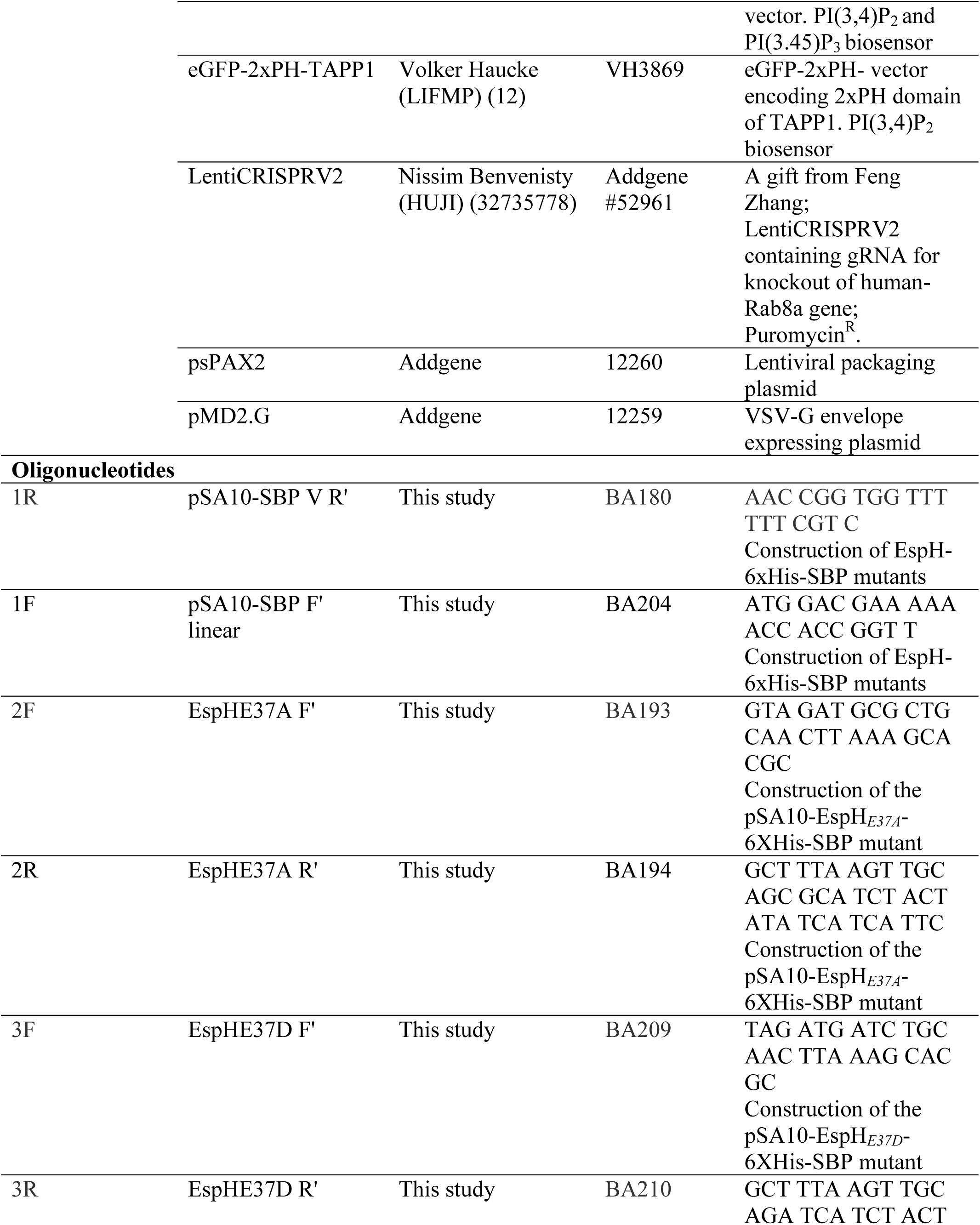
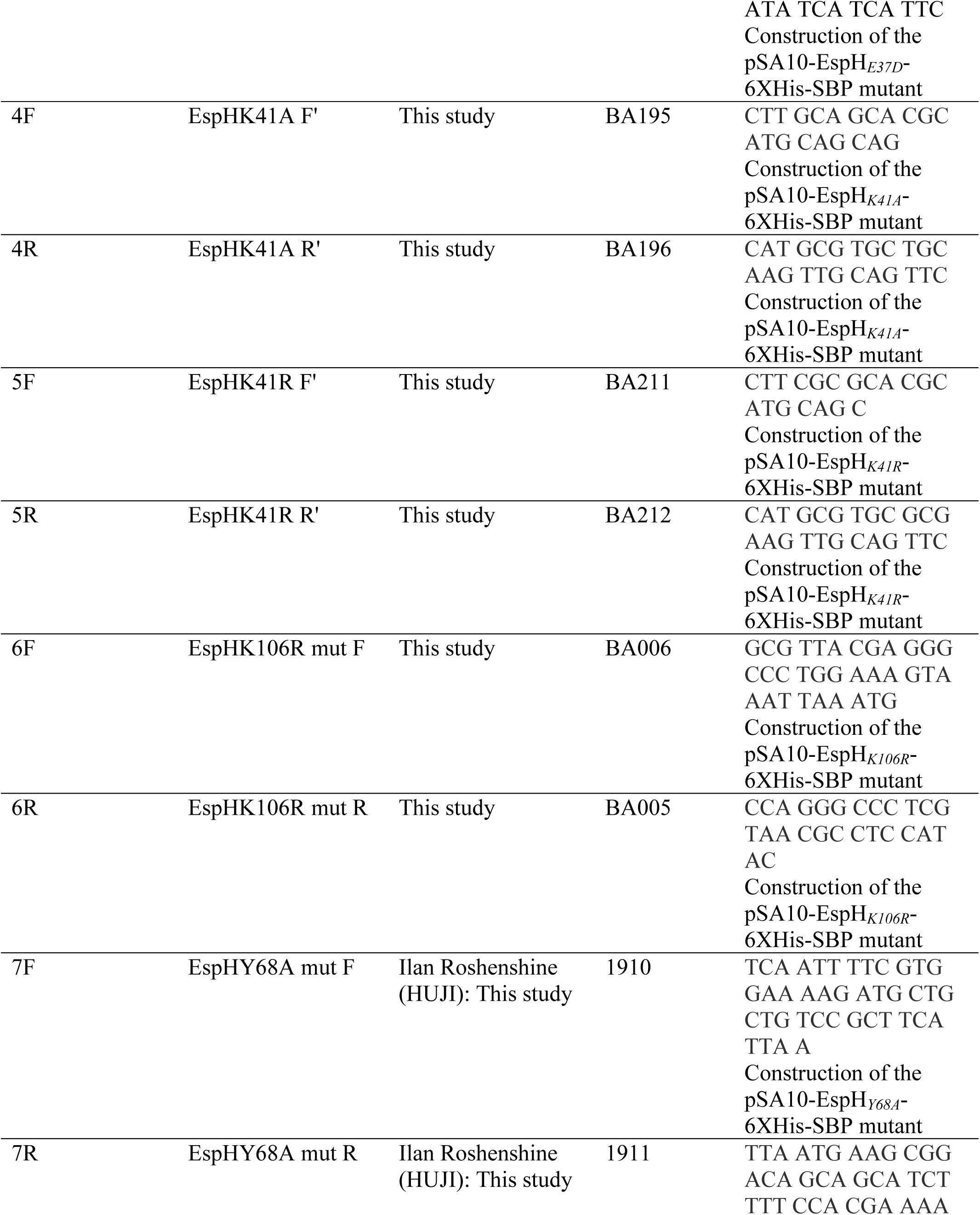
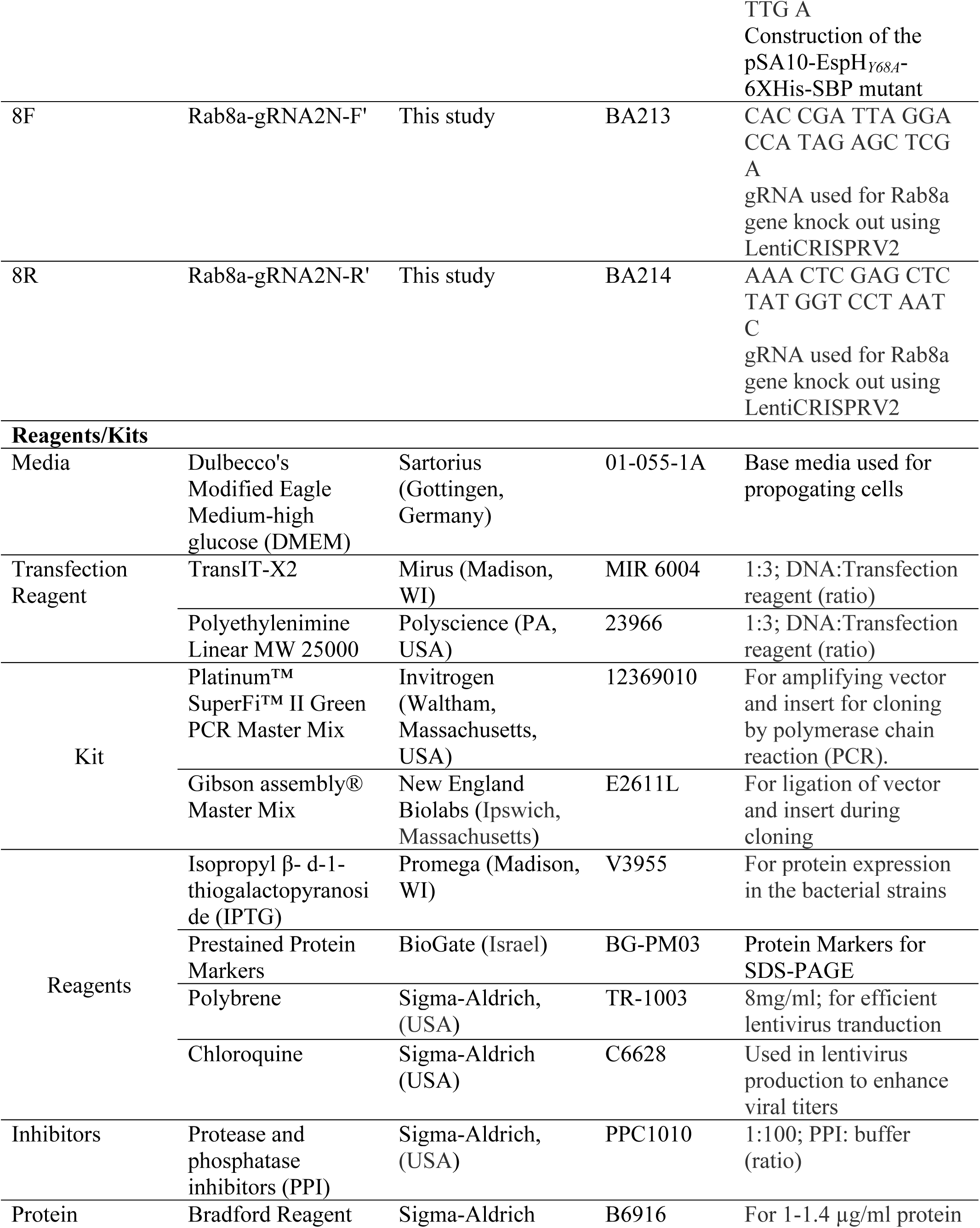
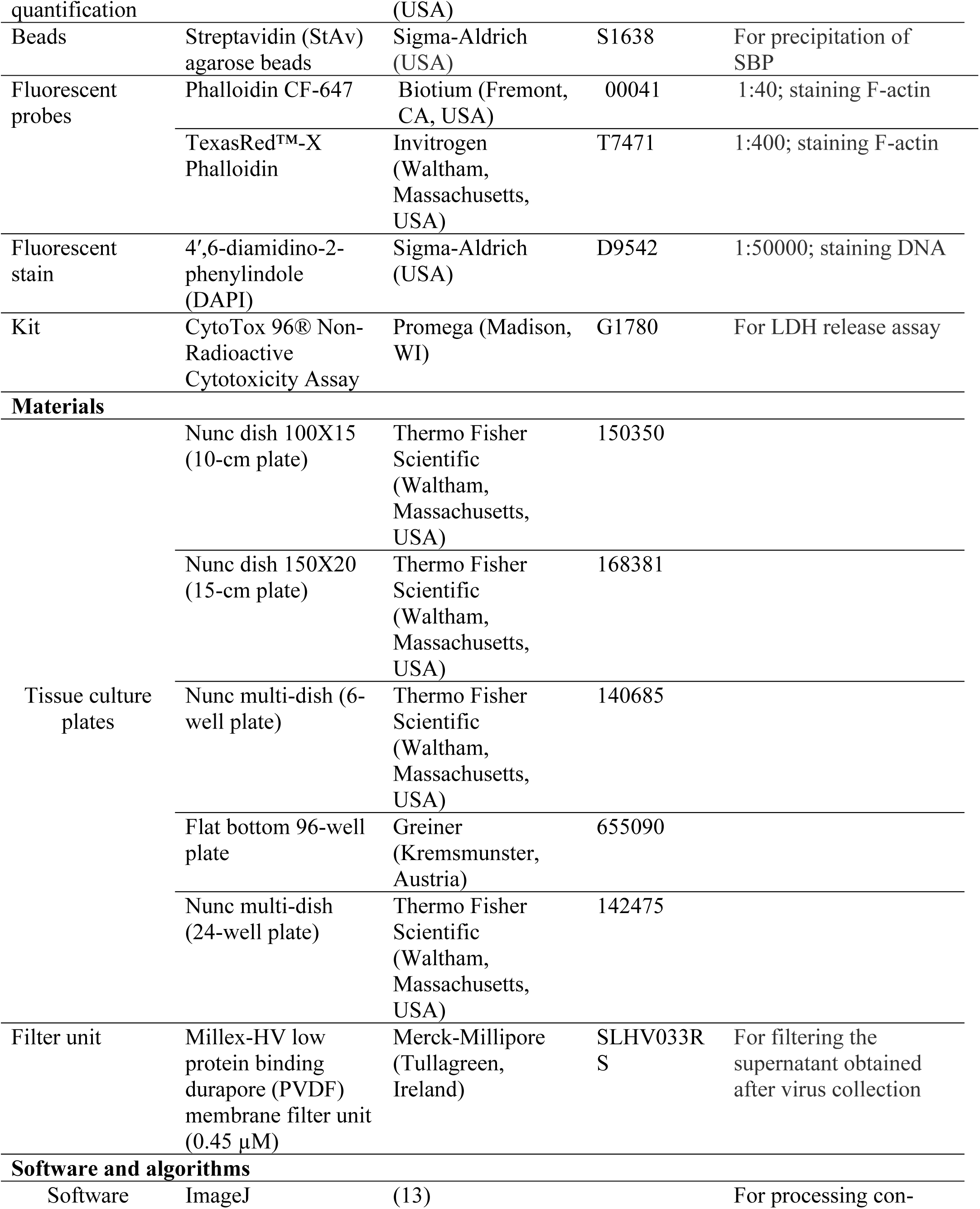
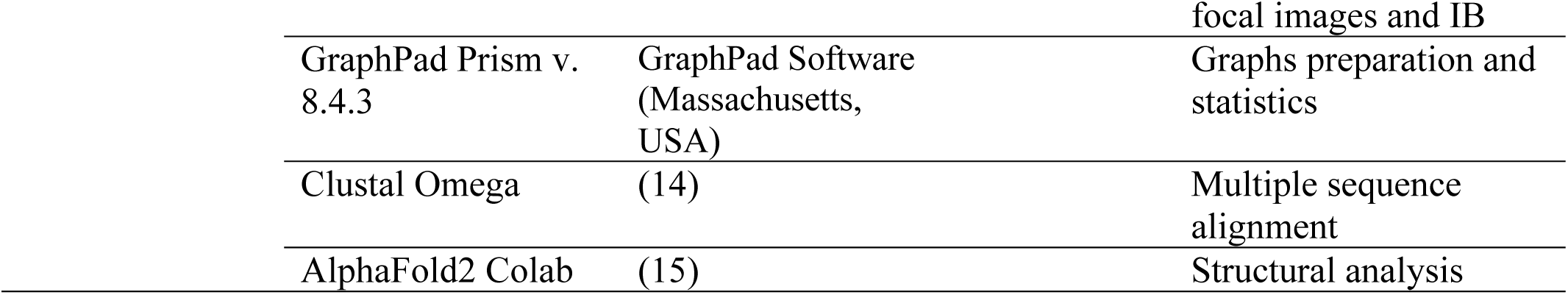

## References

Boddy KC, Zhu H, D’Costa VM, Xu C, Beyrakhova K, Cygler M, Grinstein S, Coyaud E, Laurent EMN, St-Germain J, Raught B, Brumell JH (2021) Salmonella effector SopD promotes plasma membrane scission by inhibiting Rab10. Nat Commun 12: 4707

Campellone KG (2010) Phosphoinositides influence pathogen surfing: EPEC rights the SHIP. Cell Host Microbe 7: 1–2

Celli J, Olivier M, Finlay BB (2001) Enteropathogenic Escherichia coli mediates antiphagocytosis through the inhibition of PI 3-kinase-dependent pathways. Embo J 20: 1245–1258

Chen Y, Machner MP (2013) Targeting of the small GTPase Rab6A’ by the Legionella pneumophila effector LidA. Infect Immun 81: 2226–2235

Cheng W, Yin K, Lu D, Li B, Zhu D, Chen Y, Zhang H, Xu S, Chai J, Gu L (2012) Structural insights into a unique Legionella pneumophila effector LidA recognizing both GDP and GTP bound Rab1 in their active state. PLoS Pathog 8: e1002528

Croise P, Estay-Ahumada C, Gasman S, Ory S (2014) Rho GTPases, phosphoinositides, and actin: a tripartite framework for efficient vesicular trafficking. Small GTPases 5: e29469

Dong N, Liu L, Shao F (2010a) A bacterial effector targets host DH-PH domain RhoGEFs and antagonizes macrophage phagocytosis. The EMBO journal 29: 1363–1376

Dong N, Liu L, Shao F (2010b) A bacterial effector targets host DH-PH domain RhoGEFs and antagonizes macrophage phagocytosis. EMBO J 29: 1363–1376

Dubreuil R, Segev N (2011) Bringing host-cell takeover by pathogenic bacteria to center stage. Cell Logist 1: 120–124

El Qaidi S, Wu M, Zhu C, Hardwidge PR (2019) Salmonella, E. coli, and Citrobacter Type III Secretion System Effector Proteins that Alter Host Innate Immunity. Adv Exp Med Biol 1111: 205–218

Encarnacao M, Espada L, Escrevente C, Mateus D, Ramalho J, Michelet X, Santarino I, Hsu VW, Brenner MB, Barral DC, Vieira OV (2016) A Rab3a-dependent complex essential for lysosome positioning and plasma membrane repair. J Cell Biol 213: 631–640

Escrevente C, Bento-Lopes L, Ramalho JS, Barral DC (2021) Rab11 is required for lysosome exocytosis through the interaction with Rab3a, Sec15 and GRAB. J Cell Sci 134

Gan J, Scott NE, Newson JPM, Wibawa RR, Wong Fok Lung T, Pollock GL, Ng GZ, van Driel I, Pearson JS, Hartland EL, Giogha C (2020) The Salmonella Effector SseK3 Targets Small Rab GTPases. Front Cell Infect Microbiol 10: 419

Gillingham AK, Sinka R, Torres IL, Lilley KS, Munro S (2014) Toward a comprehensive map of the effectors of rab GTPases. Dev Cell 31: 358–373

Goody PR, Heller K, Oesterlin LK, Muller MP, Itzen A, Goody RS (2012) Reversible phosphocholination of Rab proteins by Legionella pneumophila effector proteins. EMBO J 31: 1774–1784

Goosney DL, Celli J, Kenny B, Finlay BB (1999) Enteropathogenic Escherichia coli inhibits phagocytosis. Infect Immun 67: 490–495

Haritan N, Bouman EA, Nandi I, Shtuhin-Rahav R, Zlotkin-Rivkin E, Danieli T, Melamed-Book N, Nir-Keren Y, Aroeti B (2023) Topology and function of translocated EspZ. MBio: e0075223

Herve JC, Bourmeyster N (2018) Rab GTPases, master controllers of eukaryotic trafficking. Small GTPases 9: 1–4

Hong Z, Pedersen NM, Wang L, Torgersen ML, Stenmark H, Raiborg C (2017) PtdIns3P controls mTORC1 signaling through lysosomal positioning. J Cell Biol 216: 4217–4233

Humphreys D, Singh V, Koronakis V (2016) Inhibition of WAVE Regulatory Complex Activation by a Bacterial Virulence Effector Counteracts Pathogen Phagocytosis. Cell Rep 17: 697–707

Hutagalung AH, Novick PJ (2011) Role of Rab GTPases in membrane traffic and cell physiology. Physiol Rev 91: 119–149

Ingmundson A, Delprato A, Lambright DG, Roy CR (2007) Legionella pneumophila proteins that regulate Rab1 membrane cycling. Nature 450: 365–369

Jean S, Kiger AA (2012) Coordination between RAB GTPase and phosphoinositide regulation and functions. Nat Rev Mol Cell Biol 13: 463–470

Kassa EG, Zlotkin-Rivkin E, Friedman G, Ramachandran RP, Melamed-Book N, Weiss AM, Belenky M, Reichmann D, Breuer W, Pal RR, Rosenshine I, Lapierre LA, Goldenring JR, Aroeti B (2019) Enteropathogenic Escherichia coli remodels host endosomes to promote endocytic turnover and breakdown of surface polarity. PLOS Pathogens 15: e1007851

Kaur P, Dudeja PK (2023) Pathophysiology of Enteropathogenic Escherichia coli-induced Diarrhea. Newborn 2: 102–113

Khan AR (2013) Oligomerization of rab/effector complexes in the regulation of vesicle trafficking. Prog Mol Biol Transl Sci 117: 579–614

Lan Y, Sullivan PM, Hu F (2019) SMCR8 negatively regulates AKT and MTORC1 signaling to modulate lysosome biogenesis and tissue homeostasis. Autophagy 15: 871–885

Lee YT, Senturk M, Guan Y, Wang MC (2024) Bacteria-organelle communication in physiology and disease. J Cell Biol 223

Li L, Kim E, Yuan H, Inoki K, Goraksha-Hicks P, Schiesher RL, Neufeld TP, Guan KL (2010) Regulation of mTORC1 by the Rab and Arf GTPases. J Biol Chem 285: 19705–19709

Lian H, Jiang K, Tong M, Chen Z, Liu X, Galan JE, Gao X (2021) The Salmonella effector protein SopD targets Rab8 to positively and negatively modulate the inflammatory response. Nat Microbiol 6: 658–671

Luo L, Wall AA, Yeo JC, Condon ND, Norwood SJ, Schoenwaelder S, Chen KW, Jackson S, Jenkins BJ, Hartland EL, Schroder K, Collins BM, Sweet MJ, Stow JL (2014) Rab8a interacts directly with PI3Kgamma to modulate TLR4-driven PI3K and mTOR signalling. Nat Commun 5: 4407

Machner MP, Isberg RR (2007) A bifunctional bacterial protein links GDI displacement to Rab1 activation. Science 318: 974–977

Miner MV, Rauch I (2024) Why put yourself on a pedestal? The pathogenic role of the A/E pedestal. Infect Immun: e0048923

Mousnier A, Schroeder GN, Stoneham CA, So EC, Garnett JA, Yu L, Matthews SJ, Choudhary JS, Hartland EL, Frankel G (2014) A new method to determine in vivo interactomes reveals binding of the Legionella pneumophila effector PieE to multiple rab GTPases. MBio 5

Muller MP, Peters H, Blumer J, Blankenfeldt W, Goody RS, Itzen A (2010) The Legionella effector protein DrrA AMPylates the membrane traffic regulator Rab1b. Science 329: 946–949

Murata T, Delprato A, Ingmundson A, Toomre DK, Lambright DG, Roy CR (2006) The Legionella pneumophila effector protein DrrA is a Rab1 guanine nucleotide-exchange factor. Nat Cell Biol 8: 971–977

Mutvei AP, Nagiec MJ, Hamann JC, Kim SG, Vincent CT, Blenis J (2020) Rap1-GTPases control mTORC1 activity by coordinating lysosome organization with amino acid availability. Nat Commun 11: 1416

Nataro JP, Kaper JB (1998) Diarrheagenic Escherichia coli. Clin Microbiol Rev 11: 142–201

Neunuebel MR, Chen Y, Gaspar AH, Backlund PS, Jr., Yergey A, Machner MP (2011) De-AMPylation of the small GTPase Rab1 by the pathogen Legionella pneumophila. Science 333: 453–456

Novick P (2016) Regulation of membrane traffic by Rab GEF and GAP cascades. Small GTPases 7: 252–256

Ochoa TJ, Contreras CA (2011) Enteropathogenic Escherichia coli infection in children. Current opinion in infectious diseases 24: 478–483

Pfeffer S, Aivazian D (2004) Targeting Rab GTPases to distinct membrane compartments. Nat Rev Mol Cell Biol 5: 886–896

Posor Y, Jang W, Haucke V (2022) Phosphoinositides as membrane organizers. Nat Rev Mol Cell Biol 23: 797–816

Puertollano R (2014) mTOR and lysosome regulation. F1000Prime Rep 6: 52

Ramachandran RP, Nandi I, Haritan N, Zlotkin-Rivkin E, Keren Y, Danieli T, Lebendiker M, Melamed-Book N, Breuer W, Reichmann D, Aroeti B (2022) EspH interacts with the host active Bcr related (ABR) protein to suppress RhoGTPases. Gut Microbes 14: 2130657

Ramachandran RP, Spiegel C, Keren Y, Danieli T, Melamed-Book N, Pal RR, Zlotkin-Rivkin E, Rosenshine I, Aroeti B (2020) Mitochondrial Targeting of the Enteropathogenic Escherichia coli Map Triggers Calcium Mobilization, ADAM10-MAP Kinase Signaling, and Host Cell Apoptosis. MBio 11

Ramachandran RP, Vences-Catalan F, Wiseman D, Zlotkin-Rivkin E, Shteyer E, Melamed-Book N, Rosenshine I, Levy S, Aroeti B (2018) EspH Suppresses Erk by Spatial Segregation from CD81 Tetraspanin Microdomains. Infect Immun 86

Ritchie JM, Waldor MK (2005) The locus of enterocyte effacement-encoded effector proteins all promote enterohemorrhagic Escherichia coli pathogenicity in infant rabbits. Infection and immunity 73: 1466–1474

Roxas JL, Monasky RC, Roxas BAP, Agellon AB, Mansoor A, Kaper JB, Vedantam G, Viswanathan VK (2018) Enteropathogenic Escherichia coli EspH-Mediated Rho GTPase Inhibition Results in Desmosomal Perturbations. Cell Mol Gastroenterol Hepatol 6: 163–180

Roxas JL, Ramamurthy S, Cocchi K, Rutins I, Harishankar A, Agellon A, Wilbur JS, Sylejmani G, Vedantam G, Viswanathan VK (2022) Enteropathogenic Escherichia coli regulates host-cell mitochondrial morphology. Gut Microbes 14: 2143224

Salomon D, Guo Y, Kinch LN, Grishin NV, Gardner KH, Orth K (2013) Effectors of animal and plant pathogens use a common domain to bind host phosphoinositides. Nat Commun 4: 2973

Sanchez-Garrido J, Ruano-Gallego D, Choudhary JS, Frankel G (2021) The type III secretion system effector network hypothesis. Trends Microbiol

Sason H, Milgrom M, Weiss AM, Melamed-Book N, Balla T, Grinstein S, Backert S, Rosenshine I, Aroeti B (2009) Enteropathogenic Escherichia coli subverts phosphatidylinositol 4,5-bisphosphate and phosphatidylinositol 3,4,5-trisphosphate upon epithelial cell infection. Mol Biol Cell 20: 544–555

Savitskiy S, Itzen A (2021) SopD from Salmonella specifically inactivates Rab8. Biochimica et biophysica acta Proteins and proteomics 1869: 140661

Schoebel S, Cichy AL, Goody RS, Itzen A (2011) Protein LidA from Legionella is a Rab GTPase supereffector. Proc Natl Acad Sci U S A 108: 17945–17950

Shenoy AR, Furniss RCD, Goddard PJ, Clements A (2018) Modulation of Host Cell Processes by T3SS Effectors. Curr Top Microbiol Immunol 416: 73–115

Sherwood RK, Roy CR (2013) A Rab-centric perspective of bacterial pathogen-occupied vacuoles. Cell Host Microbe 14: 256–268

Shtuhin-Rahav R, Olender A, Zlotkin-Rivkin E, Bouman EA, Danieli T, Nir-Keren Y, Weiss AM, Nandi I, Aroeti B (2023) Enteropathogenic E. coli infection co-elicits lysosomal exocytosis and lytic host cell death. MBio: e0197923

Silberger DJ, Zindl CL, Weaver CT (2017) Citrobacter rodentium: a model enteropathogen for understanding the interplay of innate and adaptive components of type 3 immunity. Mucosal immunology 10: 1108–1117

Slater SL, Sagfors AM, Pollard DJ, Ruano-Gallego D, Frankel G (2018) The Type III Secretion System of Pathogenic Escherichia coli. Curr Top Microbiol Immunol 416: 51–72

Smith K, Humphreys D, Hume PJ, Koronakis V (2010) Enteropathogenic Escherichia coli recruits the cellular inositol phosphatase SHIP2 to regulate actin-pedestal formation. Cell Host Microbe 7: 13–24

Spano S, Galan JE (2018) Taking control: Hijacking of Rab GTPases by intracellular bacterial pathogens. Small GTPases 9: 182–191

Spano S, Gao X, Hannemann S, Lara-Tejero M, Galan JE (2016) A Bacterial Pathogen Targets a Host Rab-Family GTPase Defense Pathway with a GAP. Cell Host Microbe 19: 216–226

Stein MP, Muller MP, Wandinger-Ness A (2012) Bacterial pathogens commandeer Rab GTPases to establish intracellular niches. Traffic 13: 1565–1588

Sun Y, Bilan PJ, Liu Z, Klip A (2010) Rab8A and Rab13 are activated by insulin and regulate GLUT4 translocation in muscle cells. Proc Natl Acad Sci U S A 107: 19909–19914

Tan Y, Luo ZQ (2011) Legionella pneumophila SidD is a deAMPylase that modifies Rab1. Nature 475: 506–509

Tu X, Nisan I, Yona C, Hanski E, Rosenshine I (2003) EspH, a new cytoskeleton-modulating effector of enterohaemorrhagic and enteropathogenic Escherichia coli. Mol Microbiol 47: 595–606

Vetter IR, Wittinghofer A (2001) The guanine nucleotide-binding switch in three dimensions. Science 294: 1299–1304

Viaud J, Gaits-Iacovoni F, Payrastre B (2012) Regulation of the DH-PH tandem of guanine nucleotide exchange factor for Rho GTPases by phosphoinositides. Adv Biol Regul 52: 303–314

Vieira OV (2018) Rab3a and Rab10 are regulators of lysosome exocytosis and plasma membrane repair. Small GTPases 9: 349–351

Walia V, Cuenca A, Vetter M, Insinna C, Perera S, Lu Q, Ritt DA, Semler E, Specht S, Stauffer J, Morrison DK, Lorentzen E, Westlake CJ (2019) Akt Regulates a Rab11-Effector Switch Required for Ciliogenesis. Dev Cell 50: 229–246 e227

Wall AA, Luo L, Hung Y, Tong SJ, Condon ND, Blumenthal A, Sweet MJ, Stow JL (2017) Small GTPase Rab8a-recruited Phosphatidylinositol 3-Kinase gamma Regulates Signaling and Cytokine Outputs from Endosomal Toll-like Receptors. J Biol Chem 292: 4411–4422

Wang B, He W, Prosseda PP, Li L, Kowal TJ, Alvarado JA, Wang Q, Hu Y, Sun Y (2021) OCRL regulates lysosome positioning and mTORC1 activity through SSX2IP-mediated microtubule anchoring. EMBO Rep 22: e52173

Weichhart T, Hengstschlager M, Linke M (2015) Regulation of innate immune cell function by mTOR. Nat Rev Immunol 15: 599–614

Wong AR, Clements A, Raymond B, Crepin VF, Frankel G (2012a) The interplay between the Escherichia coli Rho guanine nucleotide exchange factor effectors and the mammalian RhoGEF inhibitor EspH. MBio 3

Wong AR, Pearson JS, Bright MD, Munera D, Robinson KS, Lee SF, Frankel G, Hartland EL (2011) Enteropathogenic and enterohaemorrhagic Escherichia coli: even more subversive elements. Molecular microbiology 80: 1420–1438

Wong AR, Raymond B, Collins JW, Crepin VF, Frankel G (2012b) The enteropathogenic E. coli effector EspH promotes actin pedestal formation and elongation via WASP-interacting protein (WIP). Cellular microbiology 14: 1051–1070

Wong AR, Raymond B, Collins JW, Crepin VF, Frankel G (2012c) The enteropathogenic E. coli effector EspH promotes actin pedestal formation and elongation via WASP-interacting protein (WIP). Cell Microbiol 14: 1051–1070

Wu B, Liu DA, Guan L, Myint PK, Chin L, Dang H, Xu Y, Ren J, Li T, Yu Z, Jabban S, Mills GB, Nukpezah J, Chen YH, Furth EE, Gimotty PA, Wells RG, Weaver VM, Radhakrishnan R, Wang XW, Guo W (2023) Stiff matrix induces exosome secretion to promote tumour growth. Nat Cell Biol 25: 415–424

Zerial M, McBride H (2001) Rab proteins as membrane organizers. Nat Rev Mol Cell Biol 2: 107–117

## References

1. Shen L, Black ED, Witkowski ED, Lencer WI, Guerriero V, Schneeberger EE, Turner JR. 2006. Myosin light chain phosphorylation regulates barrier function by remodeling tight junction structure. J Cell Sci 119:2095–106.

2. Nadler C, Shifrin Y, Nov S, Kobi S, Rosenshine I. 2006. Characterization of enteropathogenic Escherichia coli mutants that fail to disrupt host cell spreading and attachment to substratum. Infect Immun 74:839–49.

3. Kassa EG, Zlotkin-Rivkin E, Friedman G, Ramachandran RP, Melamed-Book N, Weiss AM, Belenky M, Reichmann D, Breuer W, Pal RR, Rosenshine I, Lapierre LA, Goldenring JR, Aroeti B. 2019. Enteropathogenic Escherichia coli remodels host endosomes to promote endocytic turnover and breakdown of surface polarity. PLoS Pathog 15:e1007851.

4. Ramachandran RP, Vences-Catalan F, Wiseman D, Zlotkin-Rivkin E, Shteyer E, Melamed-Book N, Rosenshine I, Levy S, Aroeti B. 2018. EspH Suppresses Erk by Spatial Segregation from CD81 Tetraspanin Microdomains. Infect Immun 86.

5. Roland JT, Lapierre LA, Goldenring JR. 2009. Alternative splicing in class V myosins determines association with Rab10. J Biol Chem 284:1213–23.

6. Schlosser-Silverman E, Elgrably-Weiss M, Rosenshine I, Kohen R, Altuvia S. 2000. Characterization of Escherichia coli DNA lesions generated within J774 macrophages. J Bacteriol 182:5225–30.

7. Nachury MV, Loktev AV, Zhang Q, Westlake CJ, Peranen J, Merdes A, Slusarski DC, Scheller RH, Bazan JF, Sheffield VC, Jackson PK. 2007. A core complex of BBS proteins cooperates with the GTPase Rab8 to promote ciliary membrane biogenesis. Cell 129:1201–13.

8. Rzomp KA, Scholtes LD, Briggs BJ, Whittaker GR, Scidmore MA. 2003. Rab GTPases are recruited to chlamydial inclusions in both a species-dependent and species-independent manner. Infect Immun 71:5855–70.

9. Huang B, Hubber A, McDonough JA, Roy CR, Scidmore MA, Carlyon JA. 2010. The Anaplasma phagocytophilum-occupied vacuole selectively recruits Rab-GTPases that are predominantly associated with recycling endosomes. Cell Microbiol 12:1292–307.

10. Xu J, Fotouhi M, McPherson PS. 2015. Phosphorylation of the exchange factor DENND3 by ULK in response to starvation activates Rab12 and induces autophagy. EMBO Rep 16:709–18.

11. Sason H, Milgrom M, Weiss AM, Melamed-Book N, Balla T, Grinstein S, Backert S, Rosenshine I, Aroeti B. 2009. Enteropathogenic Escherichia coli subverts phosphatidylinositol 4,5-bisphosphate and phosphatidylinositol 3,4,5-trisphosphate upon epithelial cell infection. Mol Biol Cell 20:544–55.

12. Posor Y, Eichhorn-Gruenig M, Puchkov D, Schoneberg J, Ullrich A, Lampe A, Muller R, Zarbakhsh S, Gulluni F, Hirsch E, Krauss M, Schultz C, Schmoranzer J, Noe F, Haucke V. 2013. Spatiotemporal control of endocytosis by phosphatidylinositol-3,4-bisphosphate. Nature 499:233–7.

13. Schneider CA, Rasband WS, Eliceiri KW. 2012. NIH Image to ImageJ: 25 years of image analysis. Nat Methods 9:671–5.

14. Madeira F, Pearce M, Tivey ARN, Basutkar P, Lee J, Edbali O, Madhusoodanan N, Kolesnikov A, Lopez R. 2022. Search and sequence analysis tools services from EMBL-EBI in 2022. Nucleic Acids Res 50:W276–W279.

15. Mirdita M, Schutze K, Moriwaki Y, Heo L, Ovchinnikov S, Steinegger M. 2022. ColabFold: making protein folding accessible to all. Nat Methods 19:679–682.

